# m6a methylation orchestrates IMP1 regulation of microtubules during human neuronal differentiation

**DOI:** 10.1101/2023.09.12.557394

**Authors:** Pierre Klein, Jasmine Harley, Harry Crook, Sofia Esteban Serna, Marija Petrić Howe, Theodoros I. Roumeliotis, Jyoti S. Choudhary, Anob M Chakrabarti, Raphaëlle Luisier, Rickie Patani, Andres Ramos

## Abstract

Neuronal differentiation requires building a complex intracellular architecture, and therefore the coordinated regulation of defined sets of genes. RNA-binding proteins (RBPs) play a key role in this regulation. However, while their action on individual mRNAs has been explored in depth, the mechanisms used to coordinate expression of the gene programs shaping neuronal morphology are poorly understood. To address this, we analysed how the paradigmatic RBP IMP1 (IGF2BP1), an essential developmental factor, selects and regulates its RNA targets during the differentiation of human neurons. We performed a combination of system-wide and molecular analyses, revealing that IMP1 developmentally transitions to and directly regulates the expression of mRNAs encoding essential regulators of the microtubule network, a key component of neuronal morphology. Furthermore, we showed that m6A methylation drives the selection of specific IMP1 mRNA targets and their protein expression during the developmental transition from neural precursors to neurons, providing a molecular principle for the onset of target selectivity.

## Introduction

During development, neurons establish inter-cellular networks that acquire, retain and respond to information in a spatiotemporally regulated manner. The architectural development of neurites and synapses, and the underlying cytoskeletal changes, require the regulation of gene expression programs by neuronal RNA-binding proteins (RBPs). While the action of these proteins on individual targets has been studied in detail, our understanding of how RBPs regulate networks of genes and orchestrate cellular processes during neuronal differentiation remains limited.

IMP1 is a well-studied RBP that regulates the localisation, stability and translation of mRNAs ^1^ and is essential for embryonic development ^2^. In developing neurons, IMP1 plays a key role in establishing neurite outgrowth and synaptogenesis ^3–5^. Molecular studies have focused on its physical and functional interactions with a small number of mRNA targets. In particular, the interaction between IMP1 and beta-actin mRNA has been used as a model system to explore the concept of an RNA ‘zipcode’ in the regulation of mRNA transport and translation in the cell ^6–8^. IMP1 transcriptome-wide binding and motif analyses have previously been performed, but only in highly proliferative cells including cancer and pluripotent human cells ^9,10^. These studies have highlighted a role of IMP1 in cellular adhesion, proliferation and survival. However, the mechanism by which IMP1 regulates global gene networks underlying the establishment of neuronal architecture remains unclear and is a critical knowledge void in neuronal development.

Here, we examine the system-wide role of IMP1 during the differentiation of human neurons, to elucidate how this paradigmatic protein is developmentally regulated and gain insight into how it controls the expression of essential gene programs. We show that IMP1 transitions to a different set of targets during neuronal specialization. This developmental transition is regulated by an increase of m6A methylation and, in turn, underlies the regulation of protein expression. Our findings establish IMP1 as an important regulator of the microtubule network during neuronal development.

## Results

### IMP1 transitions to a new set of neuronal targets in a regulated fashion during differentiation

RBPs have cell-type specific roles, with key functional determinants considered to be the mRNAs to which they bind and the specific sites of protein-RNA interaction. IMP1 plays multiple roles that are essential to the development of the nervous system, including synaptogenesis, dendritic arborisation and axonal pathfinding, among others ^4,11^. While IMP1 interaction with ACTB mRNA underlies an important function in neuronal development, its global interactome and role in gene regulation in neurons has not yet been explored. An essential question is whether the regulation of complex morphological neurodevelopmental processes requires IMP1 to bind to a broader set of RNAs than previously recognized. Notably, transcriptome-wide data are largely limited to highly proliferating cells, where IMP1 predominantly interacts with non-neuronal pathways.

Here we have used a human induced pluripotent stem cell (hiPSC) differentiation model, and interrogated the developmental transition from neural precursor cells (NPCs) to (isogenic) neurons. In this process, differentiating NPCs undergo major morphological changes to become neurons by building a network of neurites and forming synapses (**Fig.1a,b and Extended Data Fig.1a,b**). Firstly, we confirmed the mainly cytoplasmic expression of IMP1 in these two stages of development and its presence in developing neurites and in synapses, the latter using pre- and post-synaptic markers SYT-1 and Homer-1 (**Fig.1b, and Extended Data Fig.1b-d).** Then, we sought to define the dynamic regulation of the IMP1 protein-RNA interactome during neuronal differentiation. To this end, we scaled up our model of neuronal differentiation to millions of cells, and performed IMP1 individual-nucleotide resolution UV crosslinking and immunoprecipitation (iCLIP) on both NPCs and neurons (**Fig.1a and Extended Data Fig.2a-h**) yielding an average of 3 million unique reads in NPCs and 4 million in neurons.

**Fig.1:**
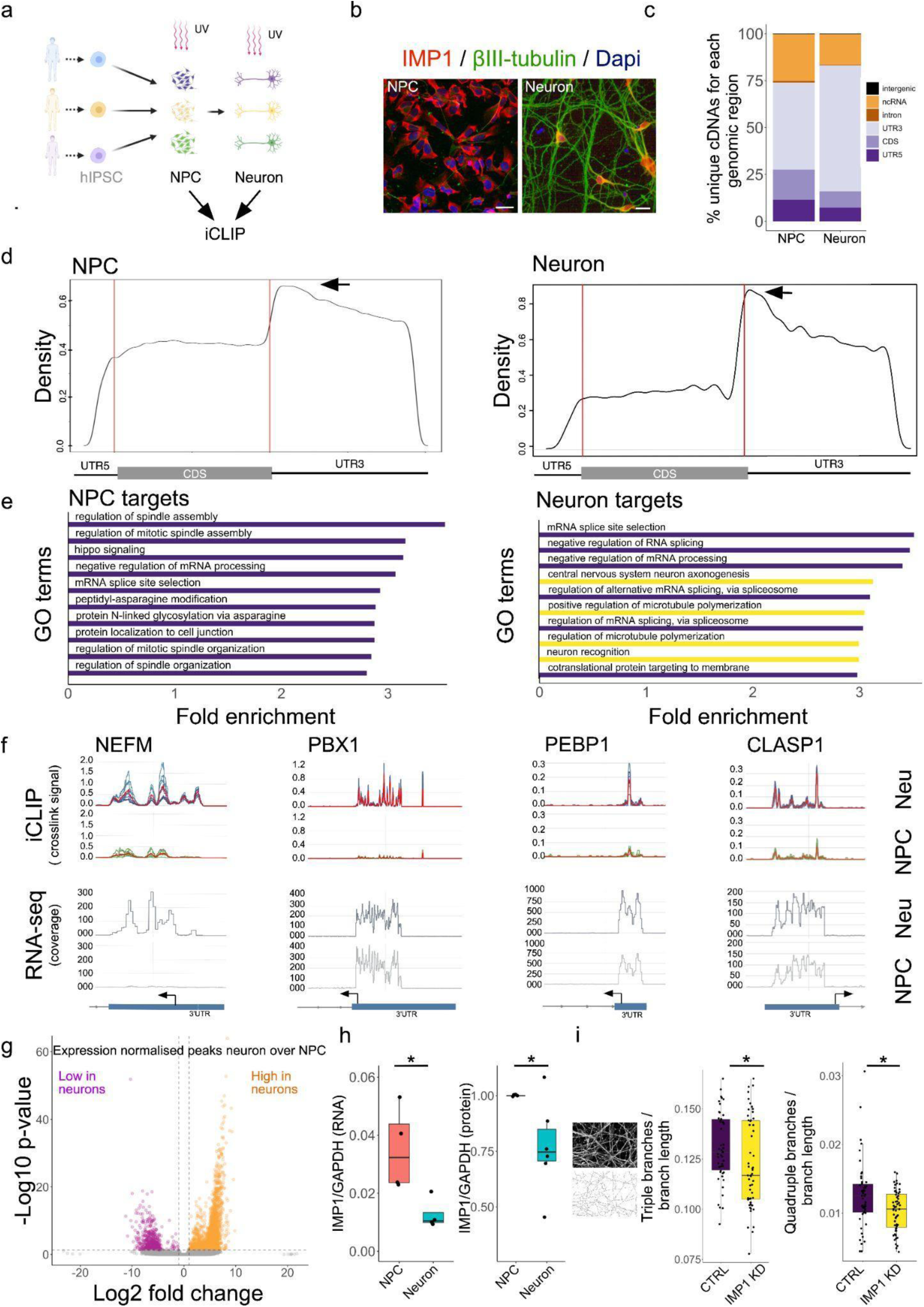
IMP1 transitions to neuronal targets and specific binding sites during differentiation. **a**, Schematic showing differentiation strategy for human neurogenesis from iPSC using a previously described protocol ^26^, that here we employed to generate the NPCs and neurons used for iCLIP. used for iCLIP. Three technical replicates from three independent iPSC lines were used for NPCs and neurons. **b**, Representative confocal images of human iPSCs differentiated into neurons immunostained for IMP1, βIII-tubulin and nuclei labelled with Dapi at NPC and neuronal stages. Scale bar, 20 μm. **c,** Barplot showing the percentage of unique cDNAs mapping to each region of the transcriptome for neurons and NPCs. Counts were normalised to the number of genomic nucleotides corresponding to each region, and percentage of total counts were calculated. Percentage is as follows: NPCs - 5’UTR 11.44, CDS 16.13, 3’UTR 46.36, Intron 0.81, ncRNA 25.13, Intergenic 0.13; MN - 5’UTR 7.32, CDS 8.71, 3’UTR 67.12, Intron 0.37, ncRNA 16.37, Intergenic 0.11. **d,** Metagene plot showing IMP1 crosslink site distribution in NPCs (left) and neurons (right). Left and right lines highlight the start and the end of the CDS respectively. Arrows indicate the peaks adjacent to the stop codons. **e**, Top GO term enrichment of genes bound by IMP1 for each cell type in neurons and NPCs. Top 10 significantly enriched terms are shown. Terms are ranked based on fold enrichment. All terms presented have a false discovery rate (FDR) <0.05. Yellow bars highlight terms related to neurogenesis or microtubules-related pathways. **f,** Mapping of IMP1 iCLIP crosslink signal (top) and RNAseq coverage signal (bottom) in neurons and NPCs. For iCLIP - tracks signal for each replicate is shown either in blue for neurons or green for NPCs and the merge of all replicates is depicted in red. The y-axis displays crosslink signal values (measured in counts per millions) for iCLIP data and read coverage for RNAseq data. The x-axis represents the genomic coordinates, specifically focusing on the last exon with the 3’ untranslated region (3’UTR). A blue rectangle highlights the location of the last exon and 3’UTR, with an arrow marking the boundary between these features with the 3’UTR side indicated, while a line-arrow indicates the intron. NEFM serves as an illustrative example of a gene displaying increased IMP1 binding in neurons, coupled with increased expression when compared to NPCs. In contrast, PBX1, PEBP1, and CLASP1 represent instances of increased IMP1 binding, yet their expression levels remain unchanged compared to NPCs. **g,** Volcano plot of IMP1 bound peaks normalised by gene expression changes in NPCs vs neurons. Preferential binding peaks in NPCs and neurons are shown respectively in pink and orange using the following criteria: log2 FC < -1 or >1 (vertical dashed line), adjusted p-value < 0.01 (horizontal dashed line). **h,** Relative expression of IMP1 over GAPDH measured by RT-qPCR (left) and WB (right) at NPC and neuronal stages. Data presented as boxplots - the center line is the median; limits are the interquartile range; and whiskers are the minimum and maximum. Data points presented are different biological replicates.For qPCR experiment n=4 independent iPSC lines. P-values calculated using a two-sided Mann–Whitney test, * P <0.05. Data presented is the mean +/-SEM. For WB experiment n= 6 technical replicates from 4 independent iPSC lines in 2 independent experiments. Neuronal values are normalised by the relative expression of the corresponding clones. Data presented is the mean +/-SEM. P-values calculated using a two-sided Mann– Whitney test, * P <0.05. **i,** Top left corner shows an example image of βIII-tubulin and skeletonization used for branching analysis. Right hand side shows quantification of triple and quadruple branches in neuronal processes based on βIII-tubulin immunostaining and normalised to branch length estimated using the βIII-tubulin signal in neurons treated with IMP1 siRNA (IMP1) vs non-targeting control (CTRL). Data presented as boxplots - the center line is the median; limits are the interquartile range; and whiskers are the minimum and maximum. Data points represent different fields of view, n=3 independent iPSC lines in 3 independent experiments. P-values calculated using two-sided Mann–Whitney test, *P < 0.05.

IMP1 crosslink sites were predominantly located in 3’UTRs, as previously reported in CLIP studies of highly-proliferative and cancer cells ^9,10^. Importantly, comparison of binding at the two stages of lineage restriction revealed that the IMP1-binding landscape is developmentally regulated, increasing its binding to the 3’UTR from 45% to 70% during the transition from NPCs to neurons (**Fig.1c, d**). We also examined whether neuronal IMP1 binding sites were enriched in the consensus motifs of IMP1 individual KH domains using a HOMER-based approach. Our analyses used a sequence of 5 bases (KH domains can recognise specifically up to 5 nucleotides). An initial analysis performed with a narrow 10-nucleotide window identified a highly enriched KH4 core consensus sequence (GGA) ^12,13^, while a follow-up analysis with a broader 30-nucleotide-window, instead, returned three motifs. The two highest scoring among these (CCGTT and ACACA) contain the core consensus sequences of KH1 (CC(or G)G) ^14^ and KH3 (CA/ACA) ^13,14^ (**Extended Data Fig.3a)**. In NPCs, although the results were less clear-cut, the KH4, KH3 and KH1 motifs were also present (data not shown). Notably, a CA sequence has been previously reported to be enriched in IMP1 target sites in previous computational studies ^10^. These results connect IMP1 neuronal target recognition with the current molecular understanding of the protein’s target specificity. Next, a comparison of mRNAs interacting with IMP1 in NPCs and neurons indicated the protein binds to a large set of targets, a substantial proportion of which are developmental stage-specific (**Fig.1e and Extended Data Fig.2e,g**). Indeed, a Gene Ontology analysis at the two developmental stages identified pathways related to early neuronal differentiation processes such as spinal cord patterning, chromatin re-organisation and synaptic development in the NPCs, while in neurons revealed pathways related to later stages of neuronal development such as axonogenesis, synapse maturation and microtubule polymerisation (**Fig.1e and Extended Data Fig.2h**).

Neuronal differentiation is accompanied by a transcriptional upregulation of a large set of genes. An important mechanistic question is whether an additional process of IMP1 target specialisation is taking place that may be important in the regulation of neuronal genes. To address this, we first compared IMP1-RNA binding iCLIP tracks of individual mRNAs, where changes during differentiation can be directly visualised (**Fig.1f and Extended Data Fig.2f**), in the context of gene expression from re-analysing RNAseq data that we previously generated (Luisier et al. 2018). This comparison highlighted that, in many cases, an increase in binding is not linked to an increase in expression. Indeed, when considering the transcripts that our iCLIP experiments identify as neuronal IMP1 targets (2567), 1218 are upregulated, 722 downregulated and 627 have a stable expression in neurons when compared to progenitors, as derived from data showed in **Extended Data Fig.2g**. Next, we explored differential binding at the transcriptome-wide level in NPCs and neurons, normalizing the changes in IMP1 peaks for RNA transcript abundance. This showed that, while transcript abundance is one important determinant of IMP1-mRNA binding in neurons, the increase in a large (1315) group of iCLIP peaks is independent of relative mRNA abundance (**Fig.1g)**, while a much smaller group of peaks (348) exhibit decreased binding. Notably, this asymmetric distribution is further accentuated when considering the peaks of mRNAs encoding proteins related to microtubule organisation and axonal development (**Extended Data Fig.3b**), confirming that the increase in IMP1 binding to these mRNAs cannot be justified only by the increase of their expression. This is consistent with the results of the Gene Ontology analysis of the developmentally regulated peaks (**Fig. 1e, Extended data Fig, 3c**), revealing an enrichment of microtubule-related mRNAs that are regulated by IMP1. Importantly, the increase in IMP1 occupancy of these targets cannot be explained by an increase in the abundance of IMP1, as our data indicate that the abundance of IMP1 in the cell decreases in the transition from progenitors to neurons, in accordance with its expression decreasing during the development of brain and other tissues ^2,15^ (**Fig.1h, Extended Data** Fig. 3d). Finally, we probed cellular protein function by examining neurite branching and synaptic development, which have previously been associated with IMP1 ^3,4^. To directly address this, we performed a morphological analysis of neurite branching, revealing a significant decrease in complexity in IMP1 knockdown (KD) neurons compared with their control counterparts (**Fig. 1i**). Furthermore, we showed that IMP1 KD decreases the size (but not the number) of SYT-1 positive synaptic puncta (**Extended Data** Fig. 3e). Together, these data show that IMP1 regulates essential neurodevelopmental processes both at the molecular and cellular levels in differentiating human neurons.

### IMP1 regulates a network of microtubule genes during human neurodevelopment

IMP1 is reported to regulate the stability, and therefore concentration, of a set of functionally related mRNAs in highly proliferating cells ^7–10,16^. However, this function is not generalisable to the global IMP1 interactome ^10^. In addition, while IMP1 has been reported to regulate the translation of ACTB mRNA by sequestering the mRNA in a folded conformation, data in highly proliferating cells indicate an association of the protein with translationally active ribosomes ^9^, and independently, that IMP1 up-regulates translation of mitochondrial mRNAs ^17^. It follows that, in principle, IMP1 could regulate the expression of sets of neuronal targets both at the protein and the RNA level. In order to determine whether IMP1 regulates the expression of sets of genes during neuronal development and whether regulation occurs at the protein or RNA level, we used mass spectrometry to profile the proteome of NPCs and neurons in control and IMP1 knockdown conditions, and intersected the data with paired RNAseq data.

In NPCs, a similar number of proteins are up and downregulated upon IMP1 KD, while in neurons twice as many proteins are downregulated compared to upregulated (**Fig.2a,b and Extended Data Fig.4a**), upon a similar KD **(Extended Data Fig.4b)**. IMP1 neuronal regulation was validated, in a representative subset of targets using quantitative immunocytochemistry and western blot (WB) as orthogonal experimental approaches (**Extended Data Fig.4c-e)**. A global functional analysis of the regulated targets indicated that IMP1 regulation of neuronal specialization is organised in gene networks underlying important cellular processes with two of the most highly represented categories of IMP1-regulated targets being related to synapses and microtubule regulation (**Fig.2c)**. The latter includes, as well as a modest group of proteins involved in myristoylation, tubulins, and regulators of microtubule stability such as kinases and essential microtubule-binding proteins. This suggests that IMP1 plays a coordinated role at multiple levels of microtubule regulation (**Fig.2c**). An unbiased analysis of the relationship between IMP1-regulated proteins revealed that they interact both physically and functionally, creating a connected network (**Extended Data Fig.4f).** This global and quantitative analysis of IMP1 regulation in neurons indicates that, in addition to to move along the microtubule system to organise local mRNA translation, as previously reported ^1,8,18^, IMP1 regulates the microtubule network itself.

**Fig.2:**
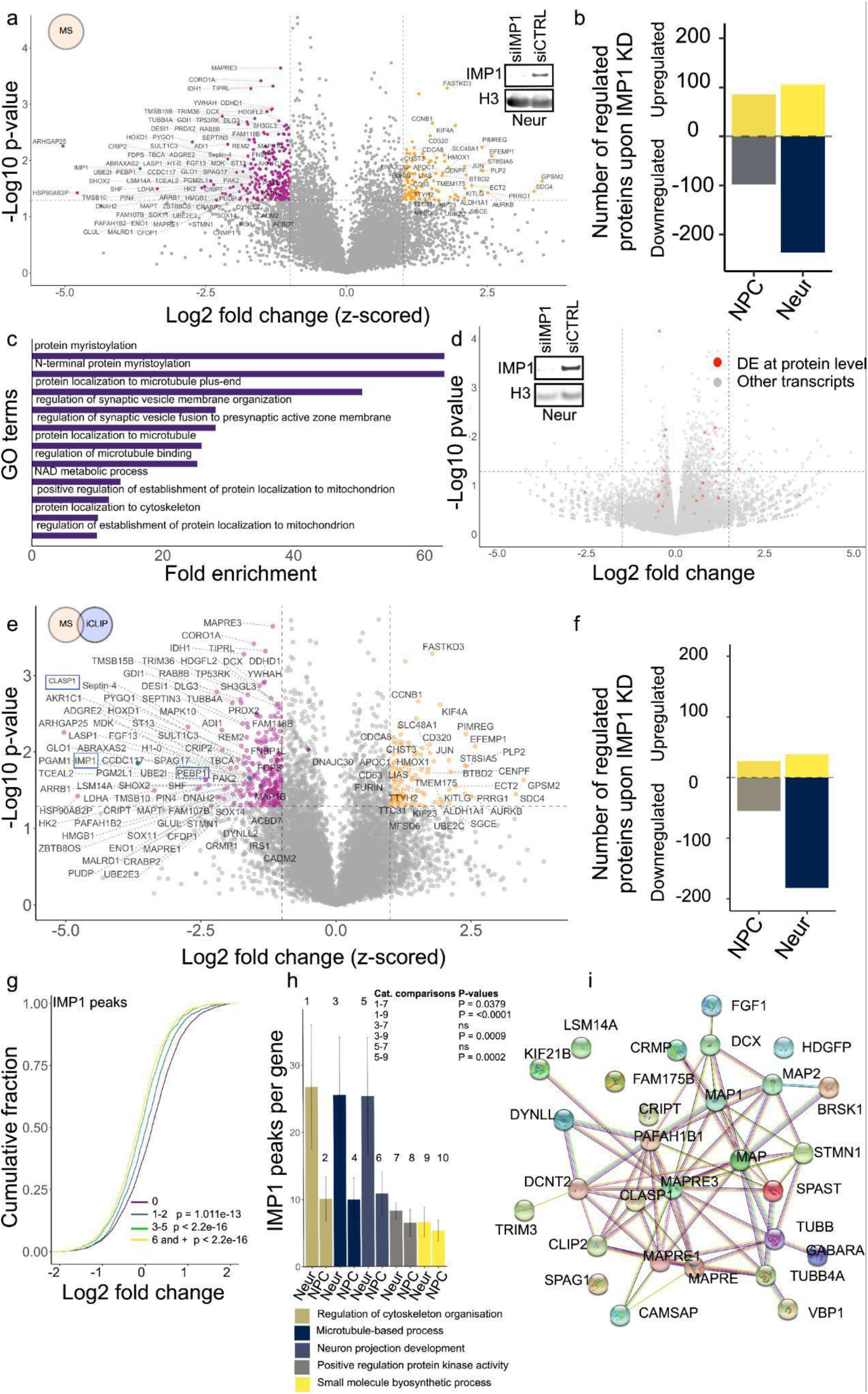
IMP1 directly regulates the network of mRNA targets. **a**, Volcano plot of differentially expressed proteins in neurons treated with IMP1 siRNA (siIMP1) vs non-targeting control (siCTRL). Downregulated, unchanged and upregulated proteins are shown respectively in pink, grey and orange with the following criteria: log2 FC (z-scored) < -1 or >1 (vertical dashed line), p-value < 0.05 (horizontal dashed line), one-sample Student’s t-test; n = 3 independent iPSC lines for each condition. Embedded representative image of Western blot showing expression of IMP1 and H3 as loading control from knockdown experiment. **b**, Barplot showing the number of proteins upregulated or downregulated in neurons and NPCs treated with IMP1 siRNA (siIMP1) vs non-targeting control (siCTRL). For downregulated proteins selection was based on log2 FC < -1, p-value < 0.05, one-sample Student’s t-test, for upregulated proteins selection was based on log2 FC > 1, p-value < 0.05, one-sample Student’s t-test. For neurons, n = 3 independent iPSC lines for each condition; for NPCs, n= 2 independent IPSC lines for each condition. **c**, Top GO term enrichments of downregulated proteins in IMP1 siRNA treated neurons (log2 FC < -1, P < 0.05). Redundancy was removed using REVIGO and top 20 significantly enriched GO terms are shown. Terms are ranked based on fold enrichment. All terms presented have a false discovery rate (FDR) <0.05. Only three proteins are found in the “protein myristoylation” and “N-terminal protein myristoylation” terms; two of these are protein phosphatases. **d**, Volcano plot of differentially expressed genes in IMP1 knockdown vs control neurons identified from the RNAseq experiment. Transcripts where corresponding proteins were also found to be differentially expressed by MS are shown in red. Vertical dashed lines indicate cut-off of log2 FC (1.5 or - 1.5), the horizontal dashed lines indicate cut-off of p-value (0.05); n = 3 independent iPSC lines for each condition. Embedded - representative image of Western blot showing expression of IMP1, and H3 as loading control from the knockdown experiment. **e**,Volcano plot of differentially expressed proteins in neurons treated with IMP1 siRNA vs non-targeting control (detected by MS) for IMP1 bound transcripts only (detected by iCLIP). Downregulated, unchanged and upregulated proteins are shown in pink, grey and orange respectively. IMP1, CLASP1 and PEBP1 are highlighted with blue dots and blue squares. For downregulated proteins selection was based on log2 FC < -1, p-value < 0.05, one-sample Student’s t-test, for upregulated proteins selection was based on log2 FC > 1, p-value < 0.05, one-sample Student’s t-test, for transcripts containing at least one IMP1 binding site. For MS, n=3 independent iPSC lines for each condition for neurons and n=2 independent iPSC lines for each condition for NPCs; for iCLIP, n=3 technical replicates from 3 independent iPSC lines for neurons, and n=6 biological+technical replicates for NPCs. **f**, Barplot showing the number of proteins upregulated or downregulated in neurons and NPCs treated with IMP1 siRNA vs non-targeting control (detected by MS) for IMP1 bound transcripts only (detected by iCLIP). For downregulated proteins selection was based on log2 FC < -1, p-value < 0.05, one-sample Student’s t-test, for upregulated proteins selection was based on log2 FC > 1, p-value < 0.05, one-sample Student’s t-test, for transcripts containing at least one IMP1 binding site. For MS, n=3 independent iPSC lines for each condition for neurons and n=2 independent iPSC lines for each condition for NPCs; for iCLIP, n=3 technical replicates from 3 independent iPSC lines for neurons, and and n=6 biological+technical replicates for NPCs. **g**, Cumulative distribution plot of log2 FC in protein expression between IMP1 KD and control in neurons detected by MS was plotted for corresponding RNA classified by the number of IMP1 peaks detected by iCLIP. For MS, n=3 independent iPSC lines for each condition; for iCLIP, n=3 technical replicates from 3 independent iPSC lines. P-values were calculated using a two-sided Kolmogorov–Smirnov test comparing 1-2 peaks, 3-5 peaks or 6 and + peaks to 0 peaks. P-values are reported on the graph. **h**, Number of IMP1 peaks per gene detected by iCLIP for GO categories related to microtubules, neuronal or other pathways classified by PANTHER - “Regulation of cytoskeleton organisation”, “Microtubule-based process”, “Neuron projection development”, “Positive regulation protein kinase activity”, “Small molecule biosynthetic process”. Number of genes in each category is in the same range (between 26 and 30). Data presented is the mean +/-SEM. Each category is represented by a number above the corresponding bar. P-values were calculated using a two-sided Mann–Whitney test and added to the plot. Comparison for the same cell type was performed. **i**, STRING analysis of protein–protein interaction network on downregulated proteins (log2 FC < -1, P < 0.05) in IMP1 siRNA treated neurons compared to control siRNA for IMP1 bound transcripts. The GO category represented is “microtubule-based process” as classified by PANTHER.

Notably, analysis of RNAseq data in both control and IMP1 KD conditions showed that only a modest number of genes are significantly up or down-regulated, and that IMP1-mediated regulation of the microtubule network occurs at the protein (rather than mRNA) level (**Fig 2d**). This is evidenced by discordant changes in IMP1 protein and RNA targets upon KD (**Fig. 2d**). Indeed, at the global level more proteins are downregulated than upregulated in neurons while this is not the case for the corresponding mRNAs. The regulation mode we observe is therefore different from the RNA stabilisation function of IMP1 reported in cancer cells and other cell lines ^7,9,10,16^.

Then, to establish whether the effect of IMP1 on the proteome is direct, we re-examined the IMP1-regulated proteome in NPCs and neurons considering only RNA targets that were directly bound by the protein using our iCLIP data. We found a striking enrichment (to a 5-fold ratio) in the proportion of downregulated compared to upregulated proteins encoded by IMP1 RNA targets upon IMP1 KD in neurons (**Fig.2e,f**), indicating that the regulation of protein expression discussed above is a direct one. Interestingly, in NPCs we also observed a modest 2-fold increase in the downregulated compared with upregulated targets upon IMP1 KD (**Fig.2f and Extended Data Fig.4g**). Beyond linking protein regulation to IMP1 binding, the changes in downregulated targets also indicate that a functional specialisation of IMP1 takes place during neuronal development.

A more in-depth analysis of the IMP1 binding pattern showed that targets positively regulated by the protein have a significantly higher number of binding sites compared to unchanged or negatively regulated targets (**Extended Data Fig.4h**). Notably, in microtubule and cytoskeleton targets, this regulation is directly linked to the number of IMP1 peaks in a transcript (**Fig.2g**). Consistently, the enrichment in IMP1 binding within the regulated targets is specific to microtubules and neuronal processes (**Fig.2h**). Together, this highlights that the regulation of microtubule assembly is a key function of IMP1 in developing neurons and confirms that this function is mediated by the direct regulation of a large gene network, as illustrated by the relationship between IMP1-regulated microtubule factors (**Fig.2i**).

### RNA methylation modulates IMP1 selection and regulation of the microtubule targets

Our data indicate that iCLIP peaks on a gene, and therefore IMP1 binding, change independently from RNA expression levels in a large set of targets (**Fig.1f,g)**, which implies an additional regulatory layer in IMP1 RNA target selection. Notably, the neuronal transcriptome has been reported to be highly enriched in m6A methylation ^19–22^. In addition, IMP1 binding to m6A has been recently reported to regulate c-Myc and cell cycle targets in cancer cells ^9^. To determine whether IMP1 regulation of the microtubule network during neuronal development is mediated by m6A methylation, we characterised the relationship between m6A methylation and IMP1 RNA binding during the NPC-to-neuron transition. While a number of recent studies have mapped m6A methylation to the transcriptome of human and mouse neurons, an m6A methylation atlas of the human transcriptome during neuronal differentiation has remained elusive. To directly compare the IMP1 RNA-binding peaks obtained from our iCLIP assays to neuronal m6A sites, we performed miCLIP on the aforementioned human NPCs and isogenic neurons (**Fig.3a, and Extended Data Fig.5a)**. The miCLIP dataset included an average of 1 million unique reads for NPCs and 4 million for neurons with high reproducibility of sites between replicates in both cell types (**Extended Data Fig.5b,c**). As expected, the crosslink sites were enriched in the evolutionarily conserved DRACH consensus motif (D = A/G/U, R = A/G, H = U/A/C) and preferentially localised around the stop codon ^19,21^ (**Fig.3b and Extended Data Fig.5d).**

**Fig.3: m6A.**
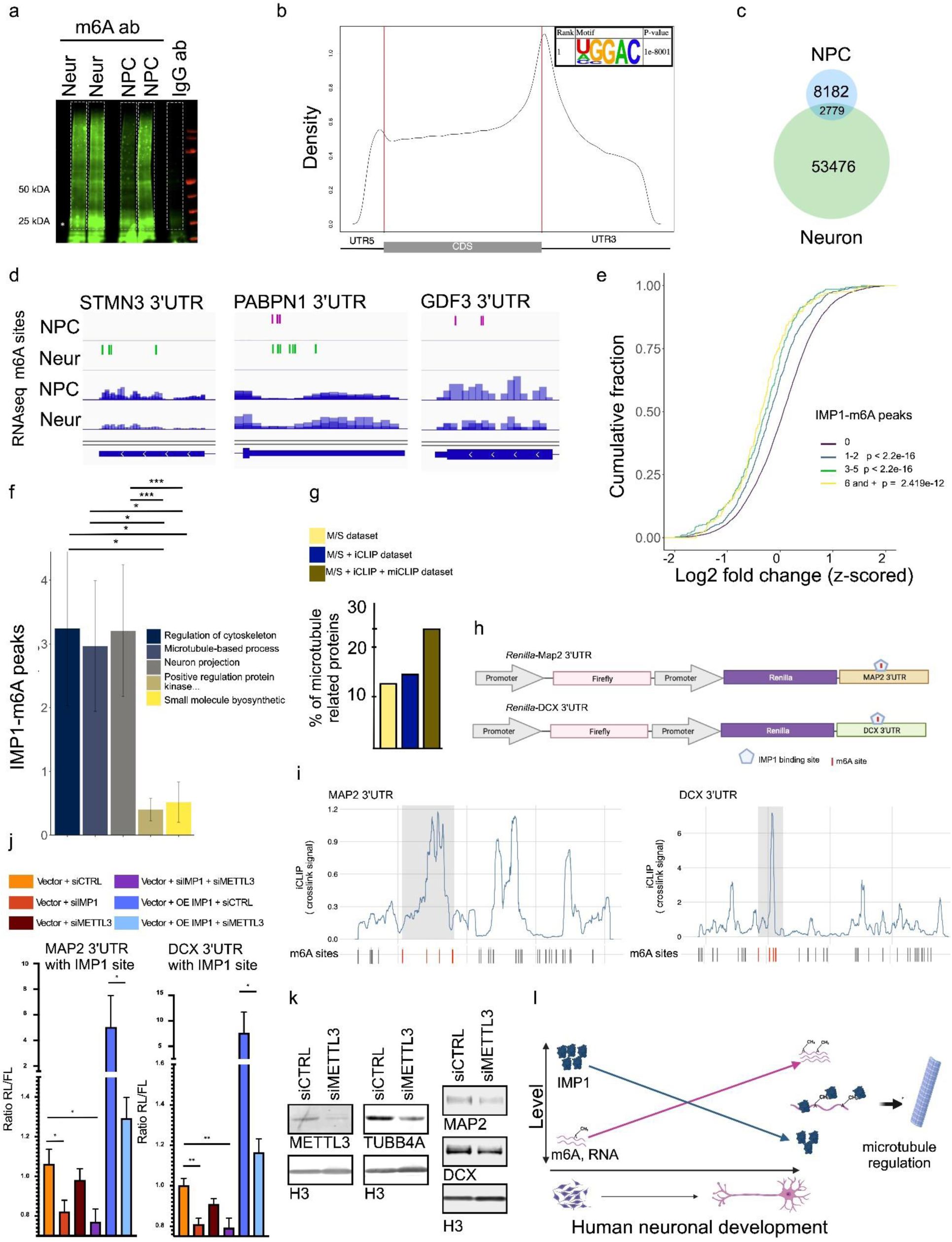
methylation modulates IMP1 binding to and regulation of microtubule targets. **a**, Representative LI-Cor scanning visualisation of nitrocellulose membrane of poly(A)+ RNA crosslinked to an m6A antibody or IgG in neurons. Visualisation uses the infrared adaptor ligated to RNA within the antibody-m6A RNA complex. The portion of the membrane excised and used to generate miCLIP libraries is shown in dashed white masks. **b**, Metagene plot showing m6A sites distribution in neurons. Embedded - consensus motif from HOMER motif discovery tools. Motif with more significant p-value is shown. **c**, Venn diagram representing the number of overlapping m6A sites between neurons and NPCs. **d**, Mapping tracks of m6A sites detected by miCLIP (top) and RNAseq coverage signal (bottom) in neurons and NPCs. The 3’UTR of three different genes is visualised using Integrative Genomics Viewer (IGV) software. **e**, Cumulative distribution plot of the log2 FC in protein expression between IMP1 knockdown and control neurons detected by MS, was plotted for corresponding RNA classified by the number of IMP1-m6A peaks detected by iCLIP and miCLIP. For MS, n=3 independent iPSC lines for each condition; for iCLIP, n=3 technical replicates from 3 independent iPSC lines, for miCLIP, n=2 technical replicates from 4 independent iPSC lines. P-values calculated using two-sided Kolmogorov–Smirnov test comparing 1-2 peaks or 3-5 peaks or 6 and + peaks to 0 peaks. P-values are reported on the plot. **f**, Number of IMP1-m6A peaks detected by overlap between miCLIP and iCLIP for GO categories related to microtubules, neuronal or other pathways classified by PANTHER - “Regulation of cytoskeleton organisation”, “Microtubule-based process”, “Neuron projection development”, “Positive regulation protein kinase activity”, “Small molecule biosynthetic process”. Number of genes in each category is in the same range (between 26 and 30). Data presented is the mean +/-SEM. For iCLIP, n=3 technical replicates from 3 independent iPSC lines, for miCLIP, n=2 technical replicates from 4 independent iPSC lines. Data presented is the mean +/-SEM. P-values calculated using a two-sided Mann– Whitney test, * P < 0.05, *** P < 0.001. **g**, Barplot of the percentage of proteins belonging to the “microtubules-based process” terms as defined by PANTHER GO classification for each of the following category : downregulated proteins as defined by MS (log2 FC < -1 and p-value >0.05), downregulated proteins with RNA targets directly bound by IMP1 as defined by MS and iCLIP, downregulated proteins with RNA targets directly bound by IMP1 through an m6A site as defined by MS, iCLIP and miCLIP. **h**, Constructs derived from the PsiCheck-2 vector that include various cloned 3’UTR sequences inserted downstream of the *Renilla* luciferase ORF. The control Firefly luciferase cassette present in the same plasmid is also shown. **i**, Mapping of IMP1 iCLIP crosslink signal (top) and m6A sites (dash) in neurons. For iCLIP, the erged signal from all replicates is shown and the y-axis displays crosslink signal values (measured in counts per millions). Only the 3’UTR portion is shown.The grey squares highlight the regions of 3’UTR used for Luciferase assay experiment, containing IMP1 binding site overlapping with m6A sites. Red dashes highlight m6A sites located in the cloned region and grey dashes all other m6A sites present in the 3’UTR. **j**, Relative luciferase activity was determined by a dual-luciferase assay system. Rluc-MAP2-UTR3 or Rluc-DCX-UTR3 constructs were co-transfected in HeLa cells with 1) siRNA CTRL, 2) siRNA CTRL + sRNA IMP1, 3) siRNA CTRL + siRNA METTL3, 4) siRNA IMP1 and siRNA METTL3, 5) siRNA CTRL with IMP1 expressing vector, or 6) siRNA METTL3 with IMP1 expressing vector. n= 6-8 replicates from at least 3 independent experiments. Data presented is the mean +/-SEM. P-values were calculated using the unpaired Student’s t-test and two-tailed Mann–Whitney test * P <0.05, ** P <0.01. Rluc stands for *Renilla* luciferase and hluc is for firefly luciferase. **k**, Western blot analysis showing expression of IMP1, METTL3, TUBB4A, DCX, MAP2 and H3 as loading control in neurons treated with non-targeting control siRNA (siCTRL) or siRNA targeting METTL3 (siMETTL3). Representative images from 2 independent experiments performed on three technical replicates from 2 independent iPSC lines. **l**, Schematic depiction of our mechanistic working model. In summary, m6A concentration increases during human neuronal differentiation, which orchestrates IMP1 regulation of microtubules.

Interestingly, starting from a similar quantity of RNA, we detected an increase in the number of m6A sites in neurons compared to NPCs (**Fig.3c)**. Consistent with this finding, we observed a concurrent increase in the level of the m6A methyl-transferase METTL3 together with the decrease in the level of the ALKBH5 demethylase (**Extended Data Fig.5f**). Importantly we found the presence of both common and specific m6A sites in NPCs and their isogenic neurons, indicating a regulated role of methylation during neuronal differentiation (**Fig.3d**). It is worth mentioning that comparing our data to 6 RNA methylation datasets from neural tissue spanning 3 species also showed a substantial overlap in m6A sites (21%) **(Extended Data Fig.5e)**. We then examined the IMP1-mediated regulation discussed in the previous paragraph in the context of methylation. In neuron-specific targets, the mRNAs encoding proteins downregulated in the IMP1 KD experiment have a higher number of IMP1-m6A peaks - where IMP1 peaks overlap with an m6A site compared to proteins that are not regulated (**Extended Data Fig.5g**). Indeed, both at the transcript and peak level we observe that IMP1-m6A overlap is higher in the regulated transcripts (**Extended Data Fig.5h**), and the overall increase in IMP1 target expression is directly correlated to the number of m6A-IMP1 sites (**Fig.3e**), with large changes observed even for single sites. In a more detailed analysis, we linked this increase to functional enrichment of IMP1 regulation, and quantified the number of IMP1-m6A sites for different GO categories. As predicted, targets related to microtubule regulation and neuronal pathways show a significantly higher number of m6A-IMP1 sites compared to unrelated pathways (**Fig.3f**). Consistent with the role that methylation plays in the overall IMP1-mediated regulation and more particularly of the microtubule network, the proportion of IMP1-upregulated proteins that are part of the microtubule network increases as we filter our MS data for i) IMP1-bound targets and ii) IMP1-m6A bound targets (**Fig.3g).** The role of m6A in IMP1-mediated gene regulation was further explored using a luciferase-based reporter assay in an established primary cell line which is more amenable to genetic manipulation. A portion of MAP2 3’UTR containing an IMP1 peak that overlapped with m6A sites was cloned downstream of *Renilla* luciferase ORF (**Fig. 3h,i**). We similarly cloned an IMP1-m6A peak in the DCX 3’UTR, to validate our observations. The luciferase activity was significantly reduced when vectors were co-transfected with an siRNA against IMP1, the m6A methyltransferase METTL3 or both. Notably, a concordant trend can be observed for both 3’UTR RNAs, where the effect of IMP1 silencing is larger than that of the silencing of the methyl-transferase. Cotransfection with both siRNAs showed a more pronounced effect than with each siRNA individually. This is consistent with an existing IMP1 binding to the specific targets can be enhanced by m6A methylation, as we proposed based on the structure and biophysical characterisation of the IMP1 KH4-m6A interaction ^23^. Importantly, overexpression of IMP1 leads to an increase of expression which is abrogated when m6A is depleted (**Fig, 3j**). The regulatory effect of m6A on IMP1 binding was validated in neurons by assessing the result of METTL3 KD on the regulated microtubules genes (**Fig.3k**). Reducing the level of METTL3 resulted in a decrease of three representative targets, including the regulators MAP2 and DCX, further confirming the role of m6A in the regulation of a neuronal network of microtubule protein expression.

## Discussion

Neuronal differentiation is coupled to an upregulation of microtubule factors that allow the assembly of cytoskeletal structures underlying the development of neurites and synapses ^24^. An important question in neuronal development is how RNA-binding proteins can control this complex process.

Here, we perform an integrated analysis of the m6A RNA methylome, transcriptome and proteome in human pluripotent stem cells during neurogenesis. These data, together with the RNA binding landscape and the functional output of the IMP1, show that this essential factor binds and regulates a large network of targets including tubulins and microtubule regulatory proteins during the critical transition between neural precursors and terminally differentiated neurons.

The understanding of how IMP1 and similar proteins coordinate morphological changes in developing neurons requires consideration of the system-wide selection of the RNA targets and regulatory action of these proteins. Our iCLIP data indicate that IMP1 binds to a large ensemble of mRNA targets in both NPCs and neurons, but also that these two ensembles are different, i.e. we observe a specialisation of IMP1 to neuronal targets upon differentiation.

IMP1 binding mediates regulation at the protein, but not the RNA level, which is different from the widespread regulation of mRNA stability function reported by many studies in highly proliferating cells ^7,9,10,16^. IMP1 regulation is dependent on cellular differentiation in the NPC- to-neuron transition, and our immunofluorescence data indicate it is necessary to build the complexity of the neurons’ network of connections. Notably, ACTB abundance was not significantly dysregulated following IMP1 knockdown, which indicates the IMP1 function we discuss here is distinct from the already-established regulation of beta-actin local translation ^1,4,25^ (**Extended Data Fig.4e**) and highlights the need for a system wide analysis of protein(s) binding and function at different stages of development.

A key question is then how IMP1 targets selectively neuronal RNAs during differentiation. Our data show a substantial increase in the transcriptome’s m6A ‘blueprint’ during the NPC- to-neuron transition in human neurodevelopment, likely mediated by the upregulation of the METTL3 m6A writer enzyme and downregulation of the ALKBH5 eraser enzyme. This change in methylation increases the number of IMP1 binding peaks in neuronal RNAs, indicating a higher occupancy on the regulated transcripts. Using first a reporter system in an established primary cell line (due to experimental tractability of genetic manipulations), and next validating these results in our human neuronal model, we confirm IMP1-regulated protein expression of microtubule targets is m6A dependent. Notably, we show that IMP1 binding and regulation of the mRNA targets is not accompanied by the upregulation of the IMP1 protein itself but rather by a decrease in the concentration of IMP1. This suggests a mechanism whereby a lower protein concentration and a site-specific increase in affinity mediated by m6A methylation together increase IMP1 selection of neuronal (microtubule) targets (**Fig.3l**). In this working model, the protein interaction with individual RNAs depends on the availability of protein and the affinity of individual binding sites. In conditions of limited protein availability, methylation enhances the functional interaction of a set of highly expressed neuronal target mRNAs with the protein. This is consistent with our recent structural and biophysical data showing that IMP1 KH4 directly recognizes m6A methylated RNA by IMP1 KH4 providing a few-fold increase in affinity and an advantage in binding ^23^.

We propose that, during neuronal differentiation, an m6A regulatory layer directs IMP1 on a subset of its neuron-specific targets in order to promote important morphological changes. Neuronal mRNAs have been reported to be highly enriched in m6A, and several non-canonical m6A readers (e.g. FMRP, hnRNPA2B1) have been reported to play a role in neuronal differentiation. The developmentally-regulated molecular mechanism of IMP1 specialisation that we describe may represent not only a key principle of IMP1-mediated differentiation, but a more widely applicable design principle in the regulation of morphological changes that accompany cell state transitions during development.

## Methods

### Ethics statement

Experimental protocols were all carried out according to approved regulations and guidelines by UCLH’s National Hospital for Neurology and Neurosurgery and UCL’s Institute of Neurology joint research ethics committee (09/0272).

### Cell culture

All cell cultures were maintained at 37°C and 5% CO_2_. The hiPSC were cultured on Geltrex coated plates with Essential 8 Medium and passaged using 0.5 mM EDTA. hiPSC-derived NPCs and neurons were cultured and differentiated according to previously published protocol ^27^. Briefly, hiPSCs were first differentiated by plating to 100% confluency in medium consisting of DMEM/F12 Glutamax, Neurobasal, L-Glutamine, N2 supplement, non-essential amino acids, B27 supplement, β-mercaptoethanol and insulin. Treatment with small molecules from day 0–7 was as follows: 1 µM Dorsomorphin, 2 µM SB431542, and 3.3 µM CHIR99021. At day 4 and 11, the cell layer was enzymatically dissociated using 1mg/ml of dispase and plated in one in two onto geltrex coated plates media containing 1 µM Rock Inhibitor. From day 8 cells were patterned for 7 days with 0.5 µM retinoic acid and 1 µM Purmorphamine. On day 14 cells were treated with 0.1 µM Purmorphamine for a further 4 days to generate spinal cord neuron precursors (NPCs). NPCs were either expanded or terminally differentiated with 0.1 µM Compound E (CE) to promote cell-cycle exit. For all experiments cells were harvested at the NPC stage or mature neurons stage (day 7 post CE treatment). Details of the lines used in this study are provided in Table 1. Two of the control lines used (control 2 and control 3) are commercially available and were purchased from Coriell (ND41866*C) and Thermo-Fisher Scientific (A18945) respectively. Information regarding the sex of the cell can be found in the key resources table. HeLa cells were cultured in DMEM supplemented with 10% FBS, 2 mM L-glutamine and 1% Penicillin/Streptomycin.

### Primary antibodies

Rabbit anti-IMP1 (MBL, RN007P), rabbit anti-IgG (Proteintech, 30000-0-AP), mouse anti-Actb (Sigma, A2228), rabbit anti-H3 (Abcam, ab201456), chicken anti-Homer-1 (Synaptic System, 160006), mouse anti-Synaptotagmin1 (Synaptic System, 105011), mouse anti-SMI-35 (Biolegend, 835603), rabbit anti-MAP1B (Proteintech, 21633-1-AP), mouse anti-MAP2 (Abcam, ab11267), chicken anti-MAP2 (Abcam, ab5392), chicken anti-beta III tubulin (Abcam, ab41489), mouse anti-beta IV tubulin (Abcam, ab11315), rabbit anti-DCX (Proteintech, 13925-1-AP), rabbit anti-MAPRE2 (Proteintech, 10364-1-AP), rabbit anti-Stathmin1 (Proteintech,11157-1-AP), mouse anti-MAPT (Abcam, ab80579), rabbit anti-CRIPT (Proteintech,11211-1-AP), mouse anti-m6A (Abcam, ab151230), rabbit anti-METTL3 (Proteintech, 15073-1-AP).

### siRNA knockdown

For RNAseq and proteomics experiments NPCs were plated in 12-well (RNAseq) or 6-well plates (proteomics) in N2B27 media at a density of 2.5 x 10^5^ cells/well or 2.5 x 10^6^ cells/well respectively. They were transfected the next day with siRNA directed against IMP1/IGF2BP1 or non-targeting siRNAs as negative control. A concentration of 30 pmol for 12-well plate and 300 pmol for 6-well plate for IMP1 was used. For m6A validation effect on protein expression, NPCs were plated in 12-well in N2B27 media at a density of 2.5 x 10^5^ cells/well. Cells were transfected the next day with 10 pmol of siRNA directed against METTL3 or non-targeting siRNAs as negative control. Lipofectamine RNAiMax was used as a transfection reagent according to the manufacturer’s instructions. After overnight incubation, the media was changed to FGF in N2B27 to maintain cells at the NPC stage or 0.1 μM Compound E in N2B27 to allow terminal differentiation to neurons. Samples were harvested for either protein or RNA extraction at 6 days after media change. Knockdown efficiency was systematically assessed by WB.

### Immunofluorescence staining

Cells were fixed in 4% paraformaldehyde in PBS for 10 min at room temperature. For permeabilization and non-specific antibody blocking, 0.3% Triton-X containing 5% bovine serum albumin (BSA) (Sigma) in PBS was added for 60 min. Primary antibodies were made up in 5% BSA and then applied overnight at 4°C. After three washes with PBS, species-specific Alexa Fluor-conjugated secondary antibody at 1:1000 dilution in 5% PBS-BSA was applied in the dark for 60 min. Cells were washed once in PBS containing Dapi, 4′,6-diamidino-2-phenylindole nuclear stain (1:1000) for 10 min. Images were taken using the Zeiss invert 880 confocal microscope or the VT-iSIM.

### Western blotting

Cells were washed with cold PBS 1X on ice and sonicated in RIPA buffer (25 mM Tris-HCl pH 7.6, 150 mM NaCl, 1% NP-40, 1% sodium deoxycholate, 0.1% SDS) supplemented with 1X Protease Inhibitor Cocktail. Supernatants were cleared of debris by 15 min centrifugation at 13000 rpm at 4°C. Protein quantification was performed using the Pierce BCA protein assay kit according to the manufacturer’s instruction. Equal quantities of proteins were supplemented with 4x Nupage loading buffer containing 1mM DTT and incubated at 90°C for 10 minutes. Samples were separated onto 4–12% Bis–Tris protein gels in 1X MES buffer and transferred onto a nitrocellulose membrane for 1 h at constant 30 V at 4°C using wet transfer. Membranes were blocked by incubation in 5% milk in PBS, for IMP1 staining for 1 h at room temperature under agitation or in 5% milk in PBS-0.05% Tween-20 (PBS-T) for all other staining. Membranes were then incubated with primary antibodies ON at 4°C in 1% milk in PBS for anti-IMP1 antibody or in 2.5% milk in PBS-T for all other antibodies either 1h at RT or ON at 4°C. Membranes were extensively washed in PBS-T and then incubated with LI-COR species-specific secondary antibodies (IRdye680 1:15000, IRdye800 1:15000) at room temperature for 1h. Unbound secondary antibody was washed in PBS-T 1X three times. Blots were imaged by Odyssey scanning (LI-COR) and quantified using Fiji software ^28^.

### RNA extraction for sequencing

IMP1 and control siRNA treated NPCs and neurons were washed with PBS 1X and harvested by centrifugation. Total RNA was extracted using Maxwell RSC simplyRNA cells kit including DNase treatment in the Maxwell RSC instrument following manufacturer’s instructions. RNA concentration and the 260/280 ratio were assessed using Nanodrop, and the Agilent 2400 Bioanalyser was used to assess quality. RNA integrity (RIN) scores were used to quality check samples.

### RNA extraction for qPCR

NPCs and neurons were washed with PBS 1X and harvested by centrifugation. Total RNA was extracted using Maxwell RSC simplyRNA cells kit including DNase treatment in the Maxwell RSC instrument following manufacturer’s instructions. RNA concentration and the 260/280 ratio were assessed using Nanodrop.

### RNA extraction for miCLIP

NPCs and neuron samples were washed with PBS 1X on ice. Content of 80 to 100% of two 6-well plates was lysed in 1.5 mL TRIzol reagent and total RNA was extracted using manufacturer’s instruction (all volumes were scaled up according to the initial volume TRIzol). RNA was resuspended in 30μl of RNase free water. RNA concentration and the 260/280 ratio were assessed using Nanodrop, and the Agilent 2400 Bioanalyser was used to assess quality. RNA integrity (RIN) scores were used to quality check samples.

### Reverse transcription and qPCR

RevertAid First Strand cDNA Synthesis kit was used to synthesise cDNA using 1 μg of total RNA and random hexamers. Appropriate dilution of the cDNA was then used in qPCR reactions containing PowerUp SYBR Green Master Mix and primer pairs, using the QuantStudio 6 Flex Real-Time PCR System. Specific amplification was determined by melt curve analysis. Gene expression levels were measured using the ΔΔCT method. IMP1 was amplified using the following pair of primers IMP1-FWD 5’-CAGGGCCGAGCAGGAAATAA - 3, IMP1-REV 5’-CAGGGATCAGGTGAGACTGC -3’ and normalised on GAPDH gene amplified with the following pair of primers: GAPDH-FWD 5’-ATGACATCAAGAAGGTGGTG-3’, GAPDH-REV 5’-CATACCAGGAAATGAGCTTG-3’ For the luciferase assay cloning vectors, 200ng of RNA from neuronal samples was reverse transcribed with superscript III reverse transcriptase (Thermo-Fisher Scientific) and Oligo(dt) using the manufacturer’s instructions.

### PolyA enrichment

Total RNA was treated with DNAse I using rigorous DNAse treatment conditions according to manufacturer’s instructions. DNAse free Poly(A)+ RNA was prepared using oligo dt dynabeads using the following protocol: 1mg of dynabeads were washed with binding buffer and resuspended in 100μl of binding buffer (20 mM Tris-HCl, pH 7.5, 1.0 M LiCl, 2 mM EDTA). Volume of RNA was adjusted to 100µl in RNAse free water and mixed with 100µl of binding buffer. Samples were incubated at 65°C for 2 min and immediately put on ice. RNA was thoroughly mixed with washed beads and tubes were rotating head over tail for 5 min at RT. Two washes with wash buffer A (10 mM Tris-HCl, pH 7.5, 0.15 M LiCl, 1 mM EDTA 10 mM Tris-HCl, pH 7.5) were performed followed by elution with 50μl of elution buffer (20 mM Tris-HCl, pH 7.5, 1 mM EDTA). Samples were then incubated at 80°C for 2 min under gentle agitation and placed on a magnetics rack. Supernatant was reused for another round of purification after beads were washed with 100µl of elution buffer and 200µl of wash buffer. RNA concentration was assessed using Nanodrop.

### Luciferase reporter assays

HeLa cells were plated on 96-well plates (Corning) and allowed to grow overnight. The next day, cells were transfected with the indicated plasmids and siRNAs using lipofectamine 2000 (Thermo-Fisher Scientific). Transfection was done with 50 ng of total plasmid (50 ng when only one plasmid was transfected or two times 25 ng for conditions with two plasmids), together with 2 pmol of total siRNA (2 pmol when only one siRNA was transfected or two times 1pmol for conditions with two siRNA) diluted in OptiMEM with 25 ng of Lipofectamine also diluted in OptiMEM. After 48 h of transfection, old media was removed and 75ul of fresh media per well was added. Luminescence of Firefly and *Renilla* luciferase were sequentially measured with a Dual-Glo® Luciferase Reporter Assay System (Promega) following the manufactured instruction using Ensight Plate Multimode Reader with default setting for luminescence measurement. For each condition, *Renilla* luciferase luminescence reads were divided by the corresponding Firefly luminescence reads. This ratio was then normalised to the average value from siRNA control conditions for each plate. For each condition at least three independent experiments in duplicates were performed. Data presented is the mean +/-SEM. Statistical analysis was performed using the unpaired Student’s t-test and the two-tailed Mann–Whitney test.

### Plasmid constructs

For luciferase assay, the dual luciferase reporter psiCHECK-2 vector (Promega) was used. 3’UTR portions were cloned downstream of Renilla luciferase gene. Vector also independently transcribes a firefly luciferase reporter which allows normalization of Renilla luciferase expression. 3’UTR fragments were generated by PCR amplification using neuronal cDNA as template, Phusion High-Fidelity PCR Master Mix with HF Buffer (NEB) following the manufacturer’s instruction and specific primers (see primer list). For the PCR, TMs were calculated using NEB’s TM calculator tool. After gel purification, PCR fragments were cloned into psiCHECK-2 using the restriction enzymes XhoI/NotI or SglI/NOTI (New England Biolabs). Ligations were performed with a quick ligation kig (New England Biolabs) and ligation products were transformed into DH5α competent *E. coli* (Thermo Fisher Scientific). Transformed DH5α were incubated overnight at 37°C on LB agar plates containing 100 μg/mL ampicillin. Colonies were inoculated overnight at 37°C in liquid cultures containing 100 μg/mL ampicillin. Plasmids were isolated using the QIAprep Spin Miniprep kit (Qiagen). The presence of the 3’UTRs was assessed using Sanger sequencing (Source Bioscience) and midipreps (Invitrogen™ PureLink™ HiPure Plasmid Filter Midiprep Kit) were performed on clones containing the correct inserts. Human IMP1 ORF cDNA clone expression plasmid was obtained from Stratech (HG18209-UT-SIB).Primers: DCX_UTR3_F 5’ AAAAACTCGAGtgcccagctgacaagactaa 3’, DCX _UTR3_R: 5’ AAAAAGCGGCCGCccatgggtggattttttctcttc, MAP2_UTR3_F: AAAAAGCGATCGCTttcattaggatggactcgt, MAP2 _UTR3_R: 5’ AAAAAGCGGCCGCttttatagctatagcttccc 3’.

### Immunofluorescence quantification

For immunofluorescence quantification a maximum projection of the images was taken from the Z stack. Using CellProfiler ^29^ the nuclei were then filtered to remove dead cells. The nuclear mask was expanded by 15 pixels and this region was defined as the cytoplasm. To define the neurites, a mask was created from βIII-tubulin, with the nuclei and cytoplasmic compartments removed. The mean intensities for each compartment were calculated, using the defined masks. For synaptic particle quantification, thresholding was used to define SYT1 particles. The particles were then quantified and the area measured. For the branching analysis, the maximum projection of βIII-tubulin underwent pre-processing to remove noise and binarization and then skeletonised using the Skeletonize3D plugin for ImageJ. The AnalyseSkeleton plugin was then used for branch quantification.

### Individual-nucleotide resolution UV-crosslinking and immunoprecipitation of protein-RNA complexes (iCLIP)

iCLIP was performed as previously described ^30^ with minor modifications. Briefly, three biological and three technical replicates for NPCs and neurons were cross-linked at 300mJ and then lysed in 1ml of IP lysis buffer. RNA fragmentation was performed with 0.4 units of RNase I and 4μl Turbo DNase I added to 1mL of protein lysate at a concentration of 1mg/mL. Optimal RNAse concentration was previously determined using a concentration gradient: low (0.4U), medium (0.8U) high (2.5 U) to 1 mg of protein lysate from NPCs and neurons. Optimal IMP1 antibody was previously determined using 1μg or 5μg of antibody to 1 mg of protein lysate. To separate protein-RNA complexes, samples were incubated with 5μg of anti-IMP1 antibody or 5μg of anti-IgG (negative control) coupled to Protein G beads at 4°C ON rotating head over tail. RNA was ligated to a pre-adenylated infrared labelled IRL3 adaptor with the following sequence:

/5rApp/AG ATC GGA AGA GCG GTT CAG AAA AAA AAA AAA /iAzideN/AA AAA AAA AAA A/3Bio/. The protein-RNA complexes were then size-separated by SDS-Page, blotted onto nitrocellulose and visualised by Odyssey scanning (LI-COR). Desired region (determined from the RNAse gradient experiment) was cut from the membrane in small pieces and RNA was released from the membrane by proteinase K digestion and incubation for 60 min at 50°C. RNA was recovered by Phenol Chloroform precipitation. cDNA was synthesised with Superscript IV Reverse Transcriptase. Reverse transcription was performed with primers containing UMIs and barcode (XXXXX) to allow multiplexing: /5Phos/ WWW *XXXXX* NNNN AGATCGGAAGAGCGTCGTGAT /iSp18/ GGATCC /iSp18/ TACTGAACCGC. cDNA molecules were purified using AMPure XP beads, then circularised using Circligase II followed by AMPure XP beads purification. After PCR amplification, libraries were size-selected by gel-purification and size distribution was assessed using Agilent 2400 Bioanalyser. Libraries were quantified by QuBit dsDNA HS Assay. Library composed of the same quantity of cDNA for each sample was sequenced as single-end 100bp reads on Illumina HiSeq 4000.

### m6A Individual-nucleotide resolution UV-crosslinking and immunoprecipitation (miCLIP)

miCLIP was performed as previously described ^31^ with minor modifications. Briefly, PolyA RNA (1μg) was brought to 30μl water and incubated with 3μl of 10X Fragmentation Buffer at 60°C for 15min. Samples were immediately placed on ice and 3.3 μl of Stop Solution was added. Samples were incubated with 10μl of 1 μg/μl of anti-m6A antibody in 500μl IP buffer (50 mM Tris-HCL, pH 7.4, 100mM NaCl, 0.05% NP-40) for 2h at 4°C rotating head over tail. RNA-anti m6A antibody complexes were crosslinked in a 6-well plate on ice twice with 0.3 J/cm-2 UV light (254 nm) in a stratalinker. Negative controls (non-crosslink and anti-IgG antibodies) were also prepared. Crosslinked RNA-anti-m6A complexes were mixed with 30μl of washed Protein G Dynabeads and incubated at 4°C for 2 hours rotating head over tail. From this point, the library preparation steps were performed as described in the iCLIP section. The correct region from the nitrocellulose membrane was cut according to previously published protocol ^32^. Libraries were sequenced with a single- end 100bp run using an Illumina Hiseq4000 platform.

### Sample preparation for proteomics analysis

Neurons and NPCs pellets were lysed in 150 μL buffer containing 1% sodium deoxycholate (SDC), 100 mM triethylammonium bicarbonate (TEAB), 10% isopropanol, 50 mM NaCl and Halt protease and phosphatase inhibitor cocktail (100X) on ice, assisted with probe sonication, followed by heating at 90°C for 5 min and re-sonicated. Protein concentration was measured with the Quick Start Bradford Protein Assay according to manufacturer’s instructions. Protein aliquots of 50 μg were reduced with 5 mM tris-2-carboxyethyl phosphine (TCEP) for 1 h at 60°C and alkylated with 10 mM iodoacetamide (IAA) for 30 min in dark, followed by overnight digestion with trypsin at final concentration 75 ng/μL (Pierce). Peptides were labelled with the TMT 10plex reagents according to manufacturer’s instructions. The mixture was acidified with 1% formic acid and the precipitated SDC was removed by centrifugation. Supernatant was then dried with a centrifugal vacuum concentrator.

### High-pH Reversed-Phase peptide fractionation of TMT labelled peptides

Peptides were fractionated with high-pH Reversed-Phase (RP) chromatography with the XBridge C18 column (2.1 x 150 mm, 3.5 μm, Waters) on a Dionex UltiMate 3000 HPLC system. Mobile phase A was 0.1% (v/v) ammonium hydroxide and mobile phase B was acetonitrile, 0.1% (v/v) ammonium hydroxide. The TMT labelled peptides were fractionated at a flow rate of 0.2 mL/min using the following gradient: 5 minutes at 5% B, for 35 min gradient to 35% B, gradient to 80% B in 5 min, isocratic for 5 minutes and re-equilibration to 5% B. Fractions were collected every 42 sec, combined orthogonally in 12 fractions and vacuum dried.

### LC-MS analysis

LC-MS analysis was performed on a Dionex UltiMate 3000 UHPLC system coupled with the Orbitrap Lumos Mass Spectrometer. Peptides were loaded onto the Acclaim PepMap 100, 100 μm × 2 cm C18, 5 μm, trapping column at 10 μL/min flow rate and analysed with the EASY-Spray C18 capillary column (75 μm × 50 cm, 2 μm) at 50°C. Mobile phase A was 0.1% formic acid and mobile phase B was 80% acetonitrile, 0.1% formic acid. For the TMT peptides, the gradient method included: 150 min gradient 5%-38% B, 10 min up to 95% B, 5 min isocratic at 95% B, re-equilibration to 5% B in 5 min and 10 min isocratic at 5% B at flow rate 300 nL/min. Survey scans were acquired in the range of 375-1,500 m/z with mass resolution of 120 k, AGC 4×10^5^ and max injection time (IT) 50 ms. Precursors were selected with the top speed mode in cycles of 3 sec and isolated for HCD fragmentation with quadrupole isolation width 0.7 Th. Collision energy was 38% with AGC 1×10^5^ and max IT 86 ms. Targeted precursors were dynamically excluded for further fragmentation for 45 seconds with 7 ppm mass tolerance.

### Immunofluorescence quantification

For immunofluorescence quantification a maximum projection of the images was taken from the Z stack. Using CellProfiler ^29^ the nuclei were then filtered to remove dead cells. The nuclear mask was expanded by 15 pixels and this region was defined as the cytoplasm. To define the neurites, a mask was created from βIII-tubulin, with the nuclei and cytoplasmic compartments removed. The mean intensities for each compartment were calculated, using the defined masks. For synaptic particle quantification, thresholding was used to define SYT1 particles. The particles were then quantified and the area measured. For the branching analysis, the maximum projection of βIII-tubulin underwent pre-processing to remove noise and binarization and then skeletonised using the Skeletonize3D plugin for ImageJ. The AnalyseSkeleton plugin was then used for branch quantification.

### Database search and protein quantification

The mass spectra were analysed in Proteome Discoverer 2.4 with the SequestHT search engine for protein identification and quantification. The precursor and fragment ion mass tolerances were set at 20 ppm and 0.02 Da respectively. Spectra were searched for fully tryptic peptides with maximum 2 missed-cleavages. TMT 6plex at N-terminus/K and Carbamidomethyl at C was selected as static modification and Oxidation of M and Deamidation of N/Q were selected as dynamic modifications.. Peptide confidence was estimated with the Percolator node and peptides were filtered at q-value<0.01 based on decoy database search. All spectra were searched against reviewed UniProt Homo Sapiens protein entries. The reporter ion quantifier node included a TMT 10plex quantification method with an integration window tolerance of 15 ppm. Only unique peptides were used for quantification, considering protein groups for peptide uniqueness. Only peptides with average reporter signal-to-noise>3 were used for protein quantification. Proteins were normalised based on total peptide signal-to-noise per sample and scaled to mean across samples (mean=100). Statistical analysis was performed in Perseus 1.6. Log2ratios of knockdowns versus matched controls were computed followed by column z-score transformation. Differentially expressed proteins were defined by one-sample t-test using the different clones as replicates. Proteins with log2 FC (z-scored) < -1 or > 1 with p-value < 0.05 were considered as differentially expressed.

### RNA-sequencing data and gene expression analysis

Poly(A)+ selected reverse stranded RNA sequencing libraries were prepared from total RNA using KAPA mRNA HyperPrep Library kit for Illumina® kit, with 200 ng of total RNA as input. Libraries were sequenced on a HiSeq 4000 platform. A total of 30 million of 100 bp-long paired-end strand-specific reads were sequenced per sample. Raw mRNA sequencing reads were trimmed using TrimGalore (https://www.bioinformatics.babraham.ac.uk/projects/trim_galore), quality evaluated using FastQC 0.11.7, (https://www.bioinformatics.babraham.ac.uk/projects/fastqc/) and aligned to GRCh38.p13 human genome using splice-aware aligner STAR v2.6.1 ^33^ with default parameters. Aligned reads were quantified using --quantMode (TranscriptomeSAM GeneCounts) function. All samples had a percentage of uniquely map reads over 90%. Differential gene expression was measured using DESeq2 ^34^ using STAR count matrix as an input (DESeqDataSetFromMatrix function). Results were generated by comparing siRNA control to siRNA IMP1 transfected cells in neurons or NPCs. A gene was considered significantly differentially expressed if p-value < 0.05 and log2 FC < -1.5 for downregulated genes and > 1.5 for upregulated genes. To remove background, genes with zero counts were excluded. For comparing gene expression level between NPCs and neurons we reanalysed previously generated RNA-seq data from the lab ^27^ - GSE98290. Fastq files from controls samples at day 14 (NPC) and day 21 (neurons) of differentiation (SRR5483813, SRR5483814, SRR5483799, SRR5483815, SRR5483812, SRR5483797, SRR5483798) were aligned to GRCh38.p13 human genome using STAR v2.6.1. and using gencode.v36.annotation.gtf file (--sjdbGTFfile). Aligned reads were quantified using -- quantMode (TranscriptomeSAM GeneCounts) function. Bedgraph files were generated using Deeptools’ bamcoverage package ^35^ with default parameters

### Processing of iCLIP data

iCLIP reads from 9 neuronal and NPCs samples and IgG control were processed according to iCLIP analysis methods using the iMaps webserver (https://imaps.genialis.com/iclip) for standardised primary analysis. iMaps is based on the icount package ^36^. Briefly, the following steps were performed: demultiplexing using experimental barcodes, UMI identification, adapter trimming, STAR pre-mapping to rRNAs and tRNAs, STAR alignment to genome, crosslink site assignment, duplicate removal by UMI sequence, peak calling (paraclu), generation of summary files based on crosslink events on gene type, biotype, or gene region. Significant crosslink sites were defined using the ‘iCount peaks’ tool on the iMaps web server while peaks were defined by clustering the significant crosslink sites using default parameters. Control quality checks (FastQC 0.11.7, https://www.bioinformatics.babraham.ac.uk/projects/fastqc/, PCR duplication ratio, quality of sequencing and alignment statistics from iMaps) were performed on each individual sample. All samples showed a high quality with low PCR duplicate ratio (1.8 to 6.1). All samples had a similar number of uniquely mapped reads (on average 7. 3E+06 for neurons and 5.7E+06 for NPC) except for two NPC samples which were further excluded in the analysis. Crosslink or peak bed files from biological and technical replicates for neurons (9) and NPCs (7) samples were merged using iMaps group function. Peaks were then called and the output bed or bedgraph files were used for further analysis. For all samples the human GRCh38 genome build and GENCODE version 36 annotation were used. For comparison between neurons and NPC sample and to account for gene expression, normalisation at gene level was performed using Deseq2 by inputting the number of IMP1 crosslink count per gene obtained from iMaps and using gene count values obtained from RNAseq experiment (GSE98290, see “RNA-sequencing data and gene expression analysis” section) as a covariate. When required normalisation for library size was performed within the DESeq2 analysis or *clipplotR* (see visualisation of iCLIP and miCLIP data method section). To obtain IMP1 expression level independent binding sites, peaks from neurons (9) and NPCs (7) samples were clustered together using the Icount clusters option from iMAPS with a window of 20 nucleotides. Resulting bed file was used as a reference and each individual sample coverage over these peaks was calculated using Bedtools map. Values from each binding site in neurons and NPCs were compared using DeSeq2 and gene count values as a covariate. Genes with less than 10 cDNA (iCLIP or RNAseq) in 5 samples were discarded using Deseq2 rowSums function. P values were calculated using a LRT test. A threshold of 1 < Log2 Fold Change < -1 and adjusted p value < 0.005 was used to determine expression independent binding sites between NPCs and neurons. PCA plots were performed using the number of crosslink counts per gene obtained from iMaps.

### miCLIP analysis and m6A site calling

miCLIP reads from 8 neuronal and NPC samples and non-crosslinked control were processed according to iCLIP analysis methods using the iMaps web server for standardised primary analysis (See “Processing of iCLIP data”). Significant crosslink sites were defined using the ‘iCount peaks’ tool on the iMaps web server while peaks were defined by clustering the significant crosslink sites using default parameters. Control quality check (FastQC report, PCR duplication ratio, quality of sequencing and alignment statistics) was performed on each individual sample. Two samples were discarded based on unique counts number and PCR duplicate ratios. The other samples showed a high quality with low PCR duplicate ratio (1.9 to 15). Correlation between replicates was assessed using DeepTools’ multibamSummary function. Default parameters were used except for binSize (50,000 bases was used).

Heatmap was then plotted with DeepTools’ plotCorrelation function using Pearson method. Mutation (CIMS) and Truncation (CITS) site calling was performed as previously described ^39^. Briefly, low-quality bases and adaptor sequences were all removed using FLEXBAR tool (-f i1.8 -as AGATCGGAAGAGCGGTTCAG --pre-trim-phred 30 -s -t sample). Reads were demultiplexed based on 5′ barcodes for individual replicates using pyCRAC ^37^. Reads were processed in pooled or separate replicate modes using the CTK package (https://zhanglab.c2b2.columbia.edu/index.php/CTK_Documentation).

In brief the barcode was stripped and added to the name of the read. Biological and technical replicates were concatenated (CIMS analysis). PCR duplicates were collapsed using pyFastqDuplicateRemover.py script from pyCRAC. The header was transformed to be compatible with CTK analysis (mawk -F ‘[_/]’ ‘/^>/{print $1”_”$2”_”$3”/”$4”#”$3”#”$2; getline($9); print $9}’). Reads were aligned to the human genome (hg38) using bwa (aln -t 4 -n 0.06 -q 20). Positions of C → T from mapped reads were obtained using the CIMS software package ^38^. Briefly, aligned reads were parsed using parseAlignment.pl to generate a bed file of read coordinates and mutation coordinates. PCR duplicates were collapsed based on read coordinated and barcode identities using tag2collapse.pl script. To get the mutations in unique tags joinWrapper.py script was used. C to T transitions were extracted and a bed file generated. Mutation reproducibility was evaluated using CIMS.pl script. For each mismatch position, the unique tag coverage (*k*) and the number of C→T transitions (*m*) were determined. C→T transitions located in DRACH motifs were called using kmer.annotate.cims.sh script based on (m ≥ 2 or m ≥ 5) and transition frequency (1% ≥ m/k ≤ 50%). Position of truncations from mapped reads were obtained using the CITS software package ^39^. Briefly, after the mutation removal step from PCR duplicates in the CIMS strategy, deletions were obtained from the mutation file using getMutationType.pl on each individual sample. Truncations events were then identified using CITS.pl script. Files with P<0.05 or *P* < 0.001 were used for further analysis. Technical and biological replicates were then concatenated. CITS and CIMS final bed files were merged using Bedtools’s mergeBed function.

### Analysis of enrichment of IMP1 peaks and IMP1-m6A peaks

IMP1 and m6A bed files were annotated using bedtools’ mergeBed package^40^ and by intersecting them with gencode.v36.annotation.gtf file downloaded from https://www.gencodegenes.org/human/release_36.html and converted into bed file using BEDOPS^41^(gtf2bed). Annotated IMP1 and m6A bed files were filtered for protein-coding genes and processed to produce a table of IMP1 or m6A counts per gene. To obtain the number of IMP1-m6A peaks per gene, an overlap between IMP1 and m6A bed files was performed using the bedtools’ intersect package. The resulting bed file was filtered for protein-coding genes and was processed to produce a table of count per genes of m6A sites bound by IMP1. The protein expression data was obtained from the proteomics dataset. The enrichment score was obtained from the Log2ratios of knockdowns versus matched controls followed by column z-score transformation. A pseudocount of 0.001 was added to the expression values to avoid division with 0. The cumulative distribution function was calculated for genes grouped by m6A count or IMP1 peaks and plotted using R (ecdf package).

### Quantification of microtubules enrichment terms

Percentage of proteins belonging to the “microtubules-based process” term” was calculated as follow: PANTHER GO analysis was performed on downregulated proteins from the MS experiment, or downregulated proteins from MS experiment containing IMP1 peaks in the corresponding RNA as defined by iCLIP, or downregulated proteins from MS experiment with m6A-IMP1 peaks in the corresponding RNA as defined by iCLIP and miCLIP. The percentage of protein belonging to the term “microtubules-based process” was quantified based on the total number of proteins in each corresponding dataset.

### Motif analysis

De novo motif enrichment was performed on peak bed files using HOMER’s (Hypergeometric Optimization of Motif EnRichment) findMotfisGenome.pl package for (i) all unique IMP1 peaks, (ii) unique IMP1 peaks for downregulated proteins - log2 FC < -1, p-value <0.05 from M/S data with the following parameters changed: search for RNA motifs (- rna), window of 10 or 30 base pair (-size 10 or 30) and a length of 5 (-rna -len 5), and (iii) IMP1-m6A bound peaks with the following parameters changed: search for RNA motifs (- rna) and a length of 5 (-rna -len 5). Default settings were used for the other parameters (http://homer.ucsd.edu/homer/motif/rnaMotifs.html).

### Gene Ontology

GO analysis was performed using protein analysis through evolutionary relationships (PANTHER) version 16.0 (http://geneontology.org) for the biological process category, with the organism set to Homo sapiens. Results were sorted hierarchically using the default parameters (test type: Fisher’s Exact, correction: FDR). When mentioned in the legend, redundant GO terms were removed using REVIGO ^42^ (Reduce & Visualize Gene Ontology) allowing medium GO term similarity.

### Protein-protein interaction network analysis

Interactions between proteins were investigated using the STRING database (Search Tool for the Retrieval of Interacting Genes/Proteins). Default parameters were used.

### Visualisation of iCLIP and miCLIP data

CLIP data were visualised in a comparative manner on individual transcripts using the software *clipplotr* (https://github.com/ulelab/clipplotr). iCLIP signals were normalised on library size and scaled to crosslinks per million. Rolling means smoothing with a sliding window of 100 nucleotides was used. When specified in the legend Integrative Genomics Viewer (IGV) software was used.

### Generation of nervous system m6A map (Meta-Analysis)

To generate a map of previously detected m6A sites in nervous tissue, bed files from published miCLIP and MeRIP-seq datasets were downloaded from Gene Expression Omnibus (GEO) repository - GSE106607, GSE106423, GSE71154, GSE29714, GSE37005 or obtained directly from publication (supplementary data) Yu et al. 2020 ^43^. When necessary, genome coordinates were converted to hg38 genome assembly using Lift Genome Annotations (https://genome.ucsc.edu/cgi-bin/hgLiftOver). Duplicated reads were first removed.

When a dataset contained information from different conditions, it was split into corresponding bed files based on sample ID. Bed files related to the nervous system (brain and neurons) were then merged using the merge command line from Bedtools v2.30.0. with default parameters. Overlap between the nervous dataset and neuron miCLIP bed files was performed using bedtool’s intersect package with default parameters.

### Metagene plots

Metagene plots were generated from CITS and CIMS bed files using MetaPlotR pipeline as previously described ^44^.

### Statistical analysis

Statistical analyses were performed using RStudio (v. 4.1.2), or GraphPad Prism 7. Normality was assessed using the D’Agostino-Pearson or Kolmogorov-Smirnov tests. For analyses where some, or all, datasets failed a normality test, a non-parametric analysis was considered. Paired or unpaired two-tailed Student’s t-test was performed when comparing two categories, except where stated otherwise. When more than two groups were compared, one-way ANOVA was used, except where stated otherwise. For cumulative distribution, a two-sided Kolmogorov–Smirnov test was used to calculate P-values. P-values less than 0.05 were considered to be significant. Graphs and error bars reflect means +/-SEM., except where stated otherwise. Information regarding statistical tests, how significance was determined and replicate number is given in figure legends.

### Data analysis

Data analyses were performed with Microsoft Excel, Graphpad Prism 7, ImageJ (v 2.3.0), RStudio (v 4.1.2) which were also used to generate graphs. Illustrations were created with BioRender.com.

## Data availability

The RNA-seq, iCLIP and miCLIP data generated for this paper have been deposited in NCBI Gene Expression Omnibus (GEO) under accession number GSE203571. RNA-seq data used to estimate the mRNA expression level in NPCs and neurons were obtained from GEO under accession number GSE98290. miCLIP and MeRIP-seq to generate nervous tissue m6A map (meta-analysis) were from GEO under accession number GSE106607, GSE106423, GSE71154, GSE29714, GSE37005 and dataset from Yu et al. 2020 ^43^. The mass spectrometry raw data have been deposited to PRIDE accession number PXD034341. All other data are available from the corresponding author upon reasonable request.

## Acknowledgements

Thanks are due to Jernej Ule and his team in particular Flora Lee, Rupert Faraway, Charlotte Capitanchik, Klara Kuret for crucial advice and discussion for the design of iCLIP and miCLIP experiments as well as the access to reagents and protocols. We would like to thank Yoh Isogai for useful discussions, help during the design of the project, and for critical reading of the manuscript. We would also like to thank Giancarlo Abis, and other members of the Ramos and Patani labs for experimental support, comments and suggestions on the project and manuscript; members of the Crick Advanced Sequencing and Light Microscopy Facilities and the Institute Cancer Research (ICR) proteomics core facility for support.

## Author contribution

P.K., R.P. and A.R. conceived the project, J.H. performed and analysed immunofluorescence experiments, H.C and P.K. performed the miCLIP experiment, T.I.R. and J.S.C. performed mass spectrometry analysis, P.K. performed all other experiments with support from MPH for cell culture work and analyses, S.E.S., A.M.C., R.L and P.K. performed bioinformatics analysis, P.K., R.P. and A.R. wrote the manuscript with input from all authors.

## Competing interest contribution

The authors declare no competing interest.

