## Supplementary Materials for "m6a methylation orchestrates IMP1 regulation of microtubules during human neuronal differentiation"

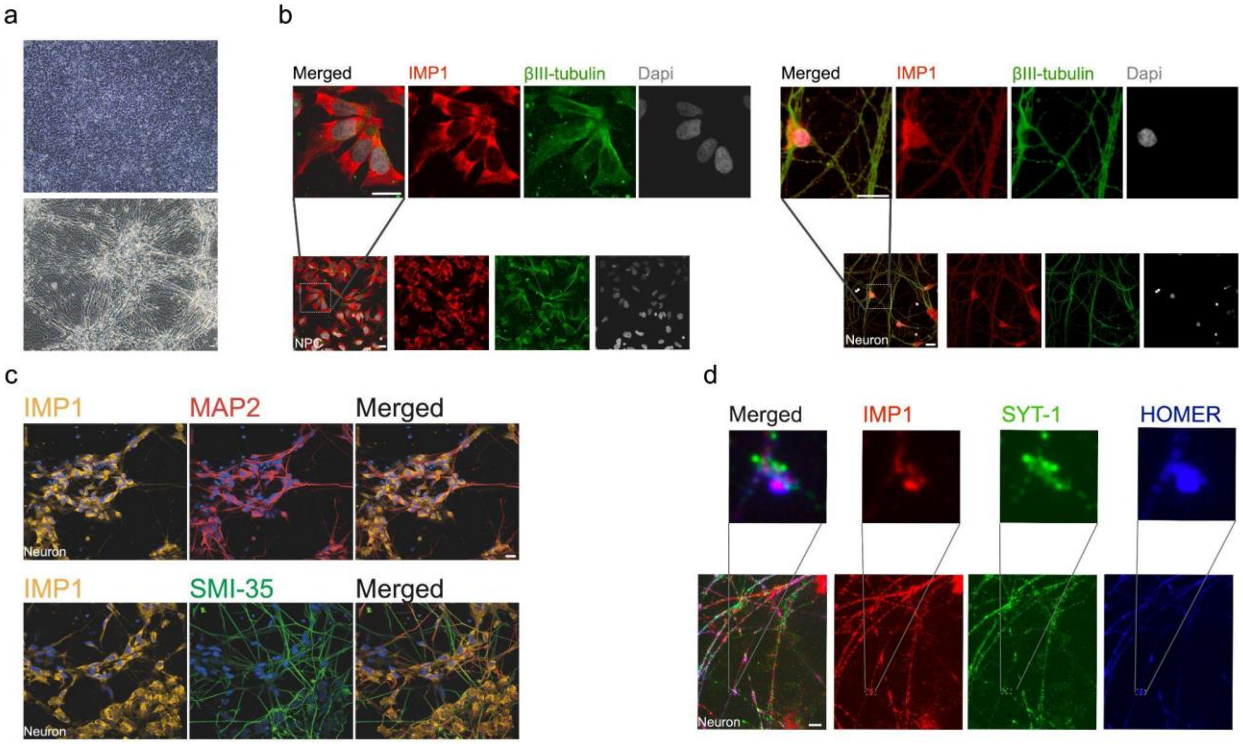


**Extended Data Fig.1 - IMP1 expression in neurites and synapses in developing neurons (related to Fig 1)**

**a**, Representative light microscopy images of neuronal precursor cells - NPC (top) and neurons (bottom) used in this study. Scale bar, 20 μm.

**b**, Representative confocal images of NPC (left) and neurons (right) immunostained for IMP1 or βIII-tubulin with nuclei labelled with Dapi. Scale bar, 20 μm.

**c**, Representative confocal images of neurons immunostained for IMP1, MAP2 and nuclei labelled with Dapi (top row); IMP1, SMI-35 and nuclei labelled with Dapi (bottom row). Scale bar, 20 μm.

**d**, Representative VT-iSIM image of neurons immunostained for IMP1, presynaptic marker SYT-1, postsynaptic marker Homer-1 (HOMER) and nuclei labelled with Dapi. Scale bar, 10 μm.

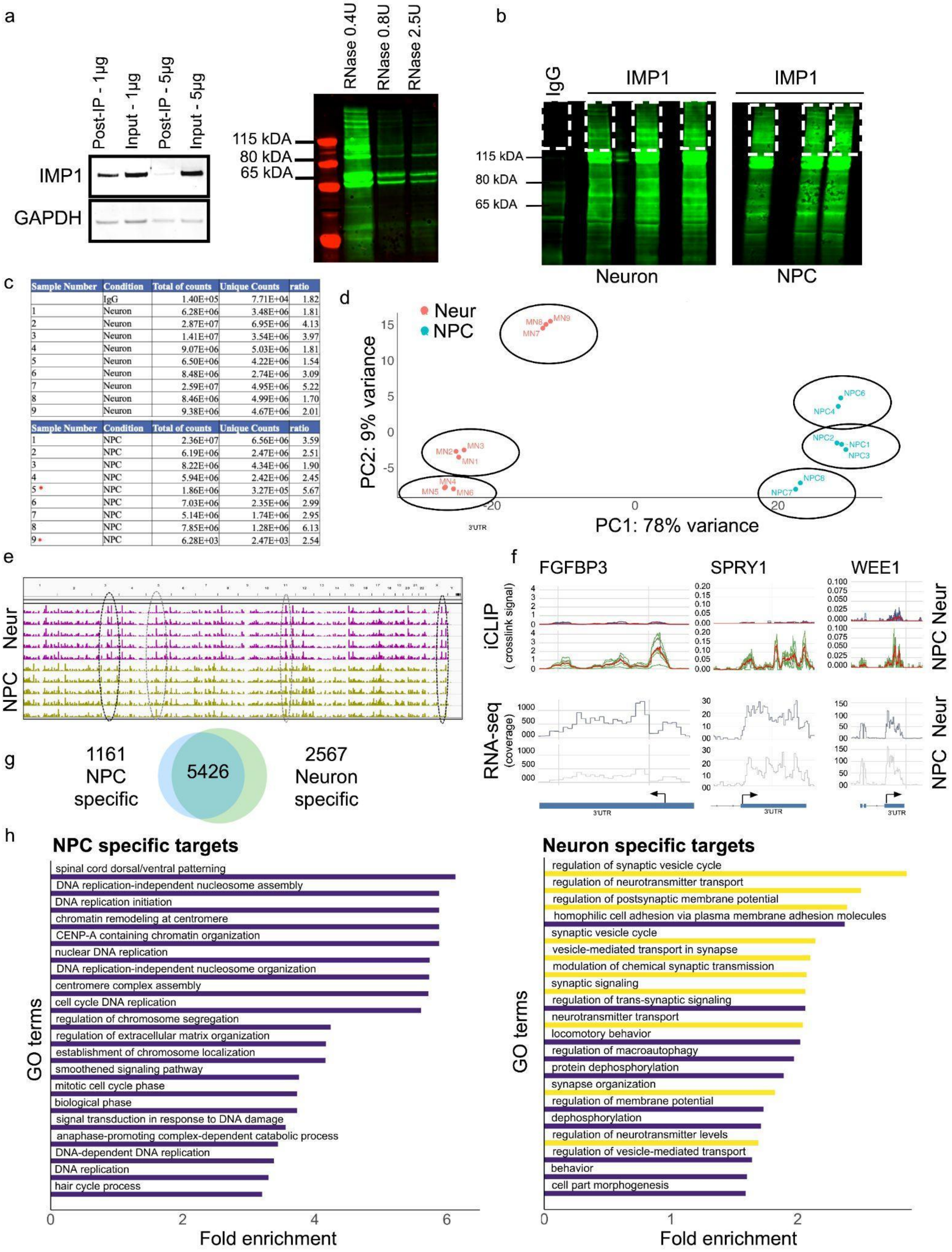


| Sample Number | Condition | Total of counts | Unique Counts | ratio |
|---|---|---|---|---|
| | IgG | 1.40E+05 | 7.71E+04 | 1.82 |
| 1 | Neuron | 6.28E+06 | 3.48E+06 | 1.81 |
| 2 | Neuron | 2.87E+07 | 6.95E+06 | 4.13 |
| 3 | Neuron | 1.41E+07 | 3.54E+06 | 3.97 |
| 4 | Neuron | 9.07E+06 | 5.03E+06 | 1.81 |
| 5 | Neuron | 6.50E+06 | 4.22E+06 | 1.54 |
| 6 | Neuron | 8.48E+06 | 2.74E+06 | 3.09 |
| 7 | Neuron | 2.59E+07 | 4.95E+06 | 5.22 |
| 8 | Neuron | 8.46E+06 | 4.99E+06 | 1.70 |
| 9 | Neuron | 9.38E+06 | 4.67E+06 | 2.01 |

| Sample Number | Condition | Total of counts | Unique Counts | ratio |
|---|---|---|---|---|
| 1 | NPC | 2.36E+07 | 6.56E+06 | 3.59 |
| 2 | NPC | 6.19E+06 | 2.47E+06 | 2.51 |
| 3 | NPC | 8.22E+06 | 4.34E+06 | 1.90 |
| 4 | NPC | 5.94E+06 | 2.42E+06 | 2.45 |
| 5 * | NPC | 1.86E+06 | 3.27E+05 | 5.67 |
| 6 | NPC | 7.03E+06 | 2.35E+06 | 2.99 |
| 7 | NPC | 5.14E+06 | 1.74E+06 | 2.95 |
| 8 | NPC | 7.85E+06 | 1.28E+06 | 6.13 |
| 9 * | NPC | 6.28E+03 | 2.47E+03 | 2.54 |

**Extended Data Fig.2 - iCLIP data capture IMP1-RNA binding in NPCs and neurons (related to Fig 1)**

**a**, Left - Western blot validation of IMP1 immunoprecipitation antibody concentration for the iCLIP experiment. A concentration of 1µg and 5µg of antibody per 1 mg of protein lysate was tested. GAPDH immunostaining was used as a loading control. Right - representative LI-Cor scanning visualisation of nitrocellulose membrane of RNA/IMP1 complexes in neurons treated with a gradient of RNAse concentrations - 0.4, 0.8 and 2.5U per 1 mL of lysate at a protein concentration of 1 mg/ml. Visualisation uses the infrared adaptor ligated to RNA.

**b**, Representative LI-Cor scanning visualisation of nitrocellulose membrane of RNA/IMP1 or RNA/IgG complexes in neurons and NPCs. Visualisation is based on the infrared adaptor ligated to RNA within protein-RNA complexes. The portion of the membrane excised and used to generate iCLIP libraries is shown in dashed white masks.

**c**, Table indicating the number of total reads, unique reads and ratio of both for each iCLIP sample. IgG condition was used as a negative technical control. Red asterisks show samples excluded from further analysis because of a significantly lower number of unique counts compared to other samples.

**d,** Principal component analysis (PCA) of cDNA counts from the iCLIP experiment of each neuron and NPC sample. Technical replicates for each clone are circled.

**e,** Mapping tracks of iCLIP crosslink signals on the whole transcriptome, visualised using Integrative Genomics Viewer software for five neuronal and five NPC samples, in pink and yellow respectively. Dotted oval highlights similar (dark grey) and dissimilar (light grey) genomic crosslink regions between NPC and neurons.

**f,** Mapping examples of IMP1 iCLIP crosslink signal (top) and RNAseq coverage signal (bottom) in neurons and NPC. For iCLIP - tracks signal for each replicate is depicted either in blue for neurons or green for NPC, while the merge of all replicates is shown in red. The y-axis displays crosslink signal values (measured in counts per millions) for iCLIP data and read coverage for RNAseq data. The x-axis represents the genomic coordinates, specifically focusing on the last exon with the 3' untranslated region (3'UTR). A blue rectangle represents the last exon and 3'UTR, with an arrow marking the boundary between these features with the 3'UTR side indicated, while a line-arrow illustrates the intron. FGFBP3 serves as an illustrative example of a gene displaying decreased IMP1 binding in neurons, coupled with increased expression when compared to NPCs. In contrast, SPRY1 and WEE1 represent instances of decreased IMP1 binding, yet their expression levels remain unchanged compared to NPCs.

**g**, Venn diagram representing the overlap between genes bound by IMP1 in neurons and NPC. When considering the expression of the 2576 neuron-specific genes, 1218 are up-regulated, 722 and down-regulated and 627 do not exhibit a significant change in expression levels.

**h,** Top GO term enrichment of genes that IMP1 uniquely binds in neurons or NPCs. Top 20 significantly enriched GO terms are shown. Terms are ranked based on fold enrichment. All terms presented have a false discovery rate (FDR) <0.05. Yellow bars highlight terms related to neurogenesis pathways.

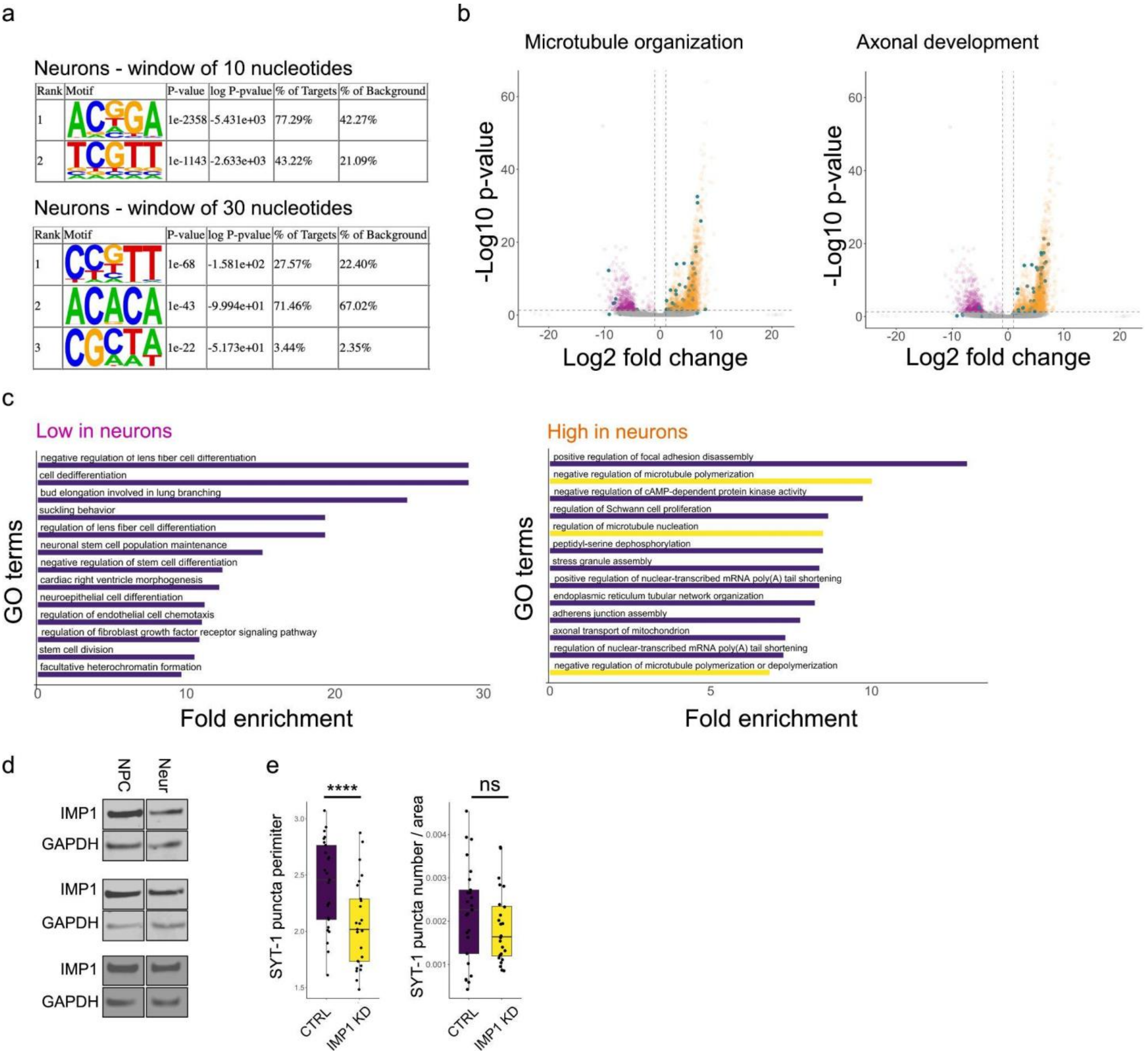


**Extended Data Fig.3 - Developmental transition of IMP1 to binding neuronal mRNAs and regulates synaptic puncta (related to Fig 1)**

**a**, Top most significantly enriched de novo sequence motifs within IMP1 binding sites in its 3'UTR targets detected by HOMER. Motif analysis performed for a window of 10 or 30 nucleotides around IMP1 binding sites. The *p*-values of the motifs, % of targets and % of background are shown.

**b**, Volcano plot of IMP1 bound peaks normalised by gene expression changes in NPC vs neurons. Peaks preferentially identified in NPC and neurons are shown respectively in pink and orange using the following criteria: log2 FC < -1 or >1 (vertical dashed lines), adjusted p-value < 0.01 (horizontal dashed line). Blue dots highlight peaks belonging to microtubule organization (GOs: regulation of cytoskeleton organization, microtubule-based process) or axonal development (GOs: neuron projection development, modulation of chemical synaptic transmission) genes. The ensemble of high-in-neuron (orange) peaks is 3.3 times larger than that in high-in-NPC (pink) peaks. When we consider only axonal development terms the ratio

is 4.5, while when microtubule organisation terms are considered, the ratio is instead of 8.1. Please note some of the blue dots are masked by orange dots.
**c**, Top GO term enrichment of IMP1 bound genes normalized by gene expression changes in NPC vs neurons - related to Figure 1g. Terms are ranked based on fold enrichment. All terms presented have a false discovery rate (FDR) $<0.05$. Yellow bars highlight terms related to microtubule regulation.
**d**, Representative Western blot images showing expression of IMP1 and GAPDH as loading control in NPC and neurons used for quantification in Fig1h.
**e**, Left - quantification of SYT-1 puncta perimeter (pixels) in neurons treated with IMP1 siRNA (IMP1) vs non-targeting control (CTRL). Data presented as boxplots - the center line is the median; limits are the interquartile range; and whiskers are the minimum and maximum. Data points represent different fields of view, n=2 independent iPSC lines in 2 independent experiments. P-values were calculated using two-sided Mann–Whitney test ** $P < 0.0001$, Right - quantification of the number of SYT-1 puncta per area of βIII-tubulin positive neurite treated with non-targeting control siRNA (CTRL) or siRNA targeting IMP1 (IMP1). Data points represent different fields of view. The boxplot presented shows the mean +/-SEM, n=2 independent iPSC lines in 2 independent experiments. P-values calculated using two-sided Mann–Whitney test, ns = not significant.

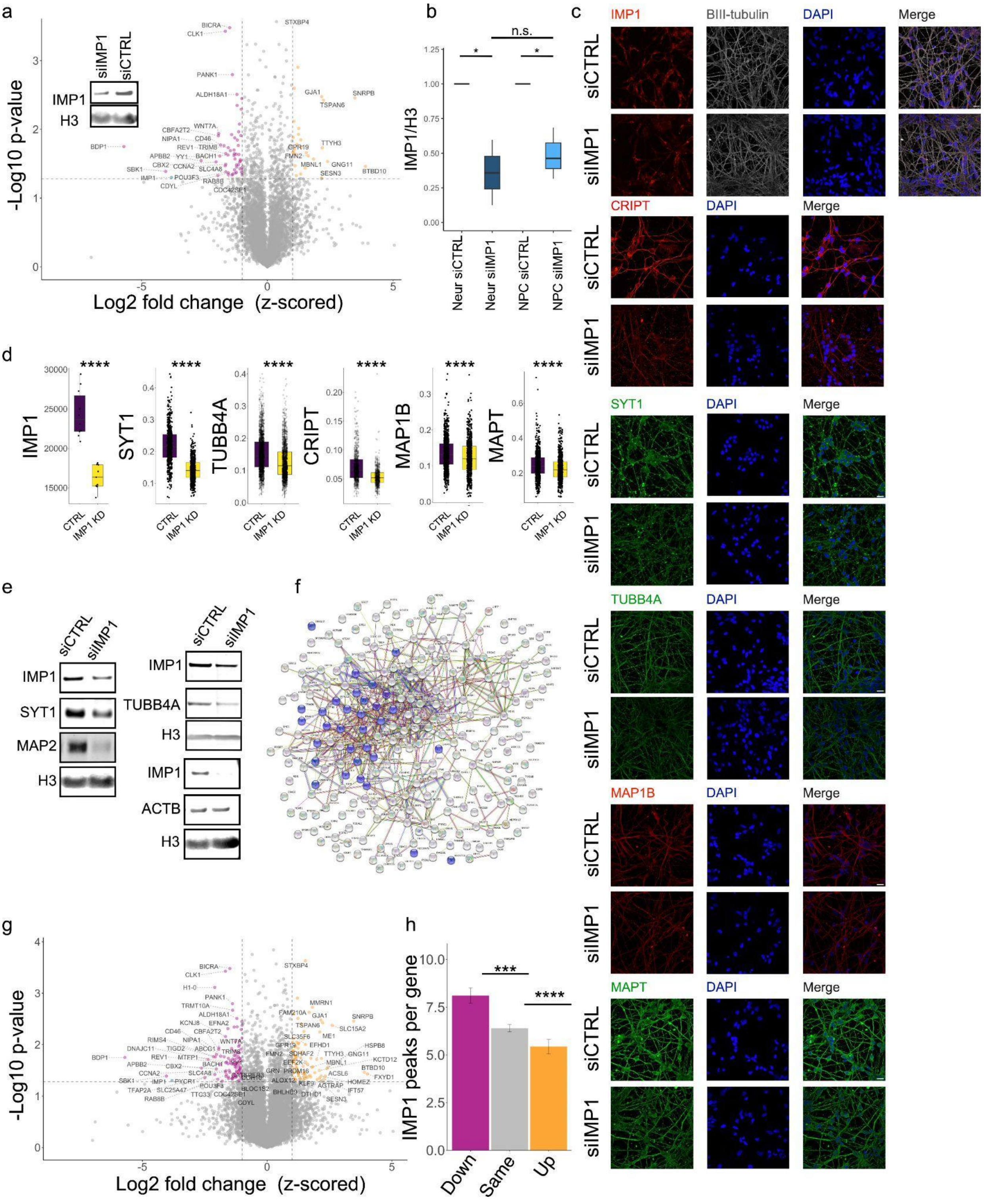


**Extended Data Fig.4 - IMP1 regulates the protein abundance resulting from bound mRNAs in NPCs and neurons (related to Fig 2)**

**a,** Volcano plot of differentially expressed proteins in NPCs treated with IMP1 siRNA (siIMP1) vs non-targeting control (siCTRL). Downregulated, unchanged and upregulated proteins are shown in pink, grey and orange respectively with the following criteria: log2 FC (z-scored) < -1 or >1 (vertical dashed lines), p-value < 0.05 (horizontal dashed line), one-sample Student's t-test; n =2 independent iPSC lines for each condition. Embedded representative image of western blot showing expression of IMP1 and H3 as loading control from the knockdown experiment.

**b**, Relative expression of IMP1 over H3 measured by Western blotting for IMP1 and control siRNA treated NPC and neuron samples sent to Mass spectrometry analysis. Boxplot presents the median (middle line), interquartile range and whiskers. n=3 independent iPSC lines, P-values calculated using a two-sided Mann–Whitney test, * P <0.05, n.s. = not significant. Data presented is the mean +/-SEM. IMP1 siRNA treated samples are normalised on relative expression of corresponding clones treated with siRNA control.

**c**, Representative confocal images of neurons treated with siRNA control or siRNA IMP1 and immunostained for IMP1, CRIPT, SYT1, TUBB4A, MAP1B or MAPT together with Dapi staining. Scale bar, 20 μm.

**d,** Quantification of IMP1, CRIPT, SYT1, TUBB4A, MAP1B, MAPT protein expression from immunostaining of neurons treated with non-targeting control siRNA (CTRL) or IMP1 siRNA (IMP1 KD). Each point represents the mean cytoplasmic intensity per field. Boxplot shows the median (middle line), interquartile range and whiskers, n=3 independent iPSC lines in 3 independent experiments. P-values calculated using two-sided Mann– Whitney test, **** P <0.0001

**e,** Representative images of Western blot showing expression of IMP1, SYT1, ACTB, MAP2, H3, IMP1, TUBB4A and H3 as loading control in neurons treated with non-targeting control siRNA (siCTRL) or siRNA targeting IMP1 (siIMP1).

**f**, STRING analysis of the protein–protein interaction network of proteins downregulated in IMP1 siRNA treated neurons (log2 FC < -1, P < 0.05). Proteins that belong to the STRING GO term "microtubule-based process" are highlighted in blue. Network protein-protein enrichment p-value < 1.0e-16.

**g**, Volcano plot of differentially expressed proteins in NPC treated with IMP1 siRNA compared to non-targeting control (detected by MS) for IMP1 bound transcripts only (detected by iCLIP). Downregulated, unchanged and upregulated proteins are shown in pink, grey and orange respectively. For downregulated proteins selection was based on log2 FC < -1, p-value < 0.05, one-sample Student's t-test, for upregulated proteins selection was based on log2 FC > 1, p-value < 0.05, one-sample Student's t-test, for transcripts containing at least one IMP1 binding site. For MS, n=2 independent iPSC lines for each condition; for iCLIP, n=6 technical + biological replicates.

**h,** Number of IMP1 peaks per gene detected by iCLIP for downregulated, unchanged or upregulated, log2 FC < -1, -1 > log2 FC < 1, log2 FC > 1 respectively - from knockdown experiments in neurons detected by MS (see Fig. 3a). Data presented is the mean +/-SEM. For MS, n=3 independent iPSC lines for each condition; for iCLIP, n=3 technical replicates from 3 independent iPSC lines. Kruskal–Wallis with Dunn's multiple comparisons test,*** P < 0.001, **** P < 0.0001.

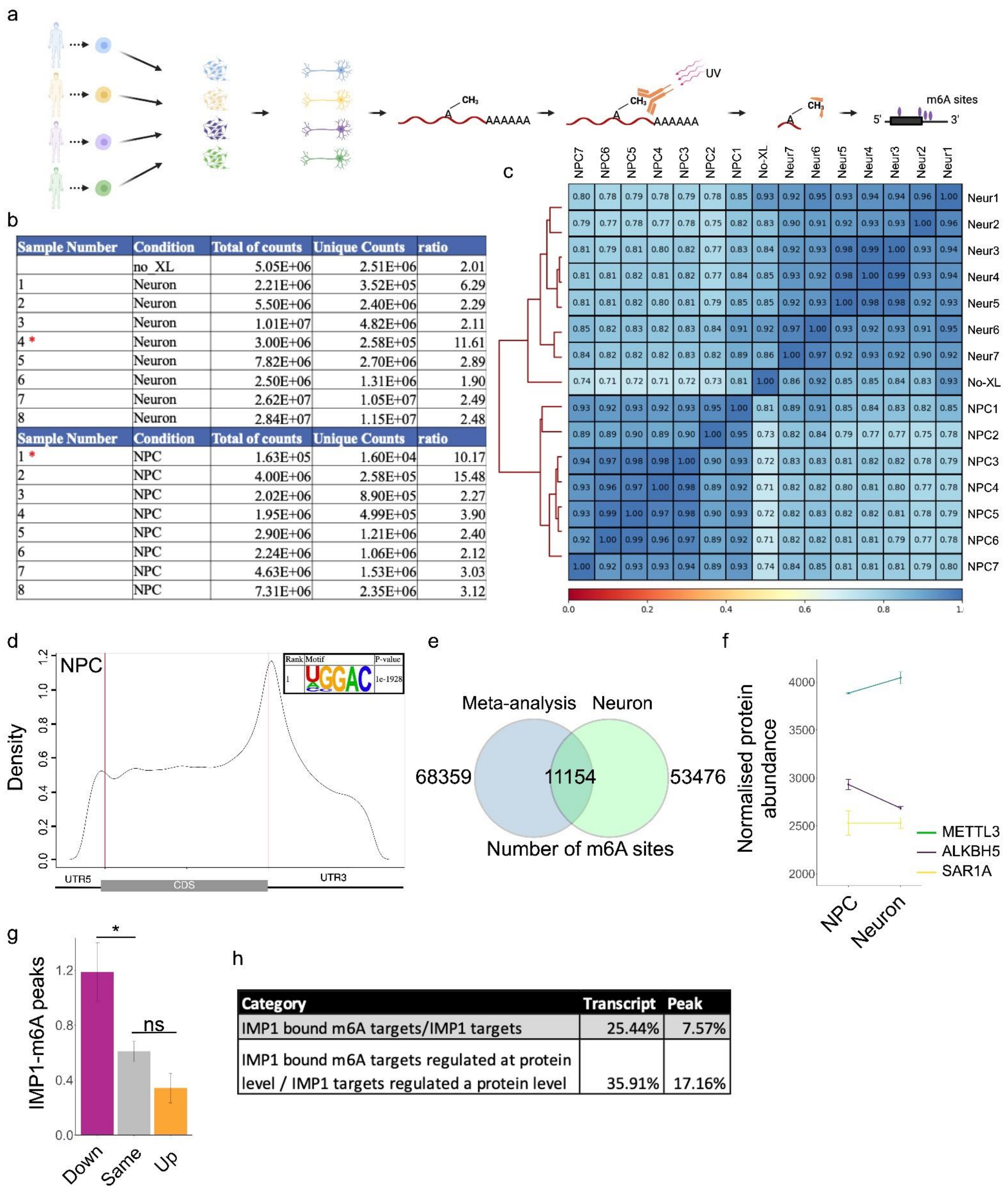


| Sample Number | Condition | Total of counts | Unique Counts | ratio |
|---|---|---|---|---|
| | no_XL | 5.05E+06 | 2.51E+06 | 2.01 |
| 1 | Neuron | 2.21E+06 | 3.52E+05 | 6.29 |
| 2 | Neuron | 5.50E+06 | 2.40E+06 | 2.29 |
| 3 | Neuron | 1.01E+07 | 4.82E+06 | 2.11 |
| 4 * | Neuron | 3.00E+06 | 2.58E+05 | 11.61 |
| 5 | Neuron | 7.82E+06 | 2.70E+06 | 2.89 |
| 6 | Neuron | 2.50E+06 | 1.31E+06 | 1.90 |
| 7 | Neuron | 2.62E+07 | 1.05E+07 | 2.49 |
| 8 | Neuron | 2.84E+07 | 1.15E+07 | 2.48 |
| **Sample Number** | **Condition** | **Total of counts** | **Unique Counts** | **ratio** |
| 1 * | NPC | 1.63E+05 | 1.60E+04 | 10.17 |
| 2 | NPC | 4.00E+06 | 2.58E+05 | 15.48 |
| 3 | NPC | 2.02E+06 | 8.90E+05 | 2.27 |
| 4 | NPC | 1.95E+06 | 4.99E+05 | 3.90 |
| 5 | NPC | 2.90E+06 | 1.21E+06 | 2.40 |
| 6 | NPC | 2.24E+06 | 1.06E+06 | 2.12 |
| 7 | NPC | 4.63E+06 | 1.53E+06 | 3.03 |
| 8 | NPC | 7.31E+06 | 2.35E+06 | 3.12 |

| Category | Transcript | Peak |
|---|---|---|
| IMP1 bound m6A targets/IMP1 targets | 25.44% | 7.57% |
| IMP1 bound m6A targets regulated at protein level / IMP1 targets regulated a protein level | 35.91% | 17.16% |

**Extended Data Fig.5 - m6A methylation and related IMP1 binding increase during neuronal development (related to Fig 3)**

**a,** Schematic showing miCLIP strategy in NPCs and neurons. Briefly, poly(A)+ RNA was extracted and UV-C crosslinked to an m6A antibody (m6A ab). RNA was then fragmented by RNase I, m6A ab-mRNA complexes were immunoprecipitated (IP) and RNA was linked to an infrared dye-labeled oligo. Complexes were run on an SDS-PAGE and transferred to a

nitrocellulose membrane. RNA was visualised using the IR dye and the region of interest was released from the membrane by proteinase K digestion. RNA was reverse transcribed, circularised, and PCR amplified. Two technical replicates from four independent iPSC lines were used for NPCs and neurons.
**b**, Table indicating the number of total reads, unique reads and ratio of both for each miCLIP sample. Non crosslinked and IgG conditions were used as negative technical controls (only the non-crosslinked condition was sequenced due to the absence of cDNA in the IgG condition). Red asterisks show samples excluded from further analysis because of a significantly lower number of unique counts and/or high ratio of unique reads over total reads compared to other samples.
**c,** Heatmap of correlation matrix for read coverage of binned genomics regions for each NPC, neuron, and non-crosslinked sample. The coloured scale depicts the correlation values which are also indicated in each square. The y-axis on the left side shows the hierarchical clustering of the samples.

**d**, Metagene plot showing m6A residue distribution in NPC. Embedded - consensus motif from HOMER motif discovery tools. Motif with the most significant p-value is shown.
**e**, Venn diagram representing the overlap between m6A sites identified in neurons and a meta-analysis of a set of published RNA methylation datasets (see method).
**f**, Abundance levels of different proteins detected by MS in NPCs and neurons treated with siRNA CTRL. Expression was normalised based on the summed abundances per sample, n=2 independent iPSC lines.
**g**, Number of IMP1-m6A peaks per transcript detected by overlapping the miCLIP and IMP1 iCLIP datasets for downregulated, unchanged or upregulated proteins - log2 FC < -1 in pink; -1 > log2 FC< 1 in grey; log2 FC > -1 in orange respectively – from IMP1 knockdown experiments in neurons. Data presented is the mean +/- SEM. For MS, n=3 independent iPSC lines for each condition, for iCLIP, n=3 technical replicates from 3 independent iPSC lines and for miCLIP, n=2 technical replicates from 4 independent iPSC lines. P-values calculated using Kruskal– Wallis with Dunn's multiple comparisons test, *P < 0.05, ns = not significant.
**h**, Table displaying 1) the proportion of targets or peaks bound by IMP1 via an m6A site, relative to the total number of targets or peaks bound by IMP1, and 2) the proportion of targets or peaks bound by IMP1 through an m6A site that are downregulated at the protein level following IMP1 knockdown, relative to all targets or peaks bound by IMP1 through m6A sites.
